## Supplementary Information for "Conformational Heterogeneity in Human Interphase Chromosome Organization Reconciles the FISH and Hi-C Paradox"

**Supplementary Note 1: Power law relation between the contact probability and mean spatial distance**

A key goal in our theory is to theoretically establish a useful relationship between the contact probabilities and the mean spatial distances between the loci. Because long chromosomes are modeled as polymers, we look to rigorous results in polymer theory for the distance distribution function, $P(r|\langle R\rangle)$ between two loci separated by $r$ with a mean distance $\langle R\rangle$. Knowledge of $P(r|\langle R\rangle)$ is needed to use Equation (3) in the main text in order to construct the Cumulative Distribution Function, $\mathrm{CDF}(R|\langle R\rangle)$ (see Supplementary Equation 4 below). There are only few polymer models for which analytic results for $P(r)$ are known.

A particularly useful result for our purposes is $P(r|\langle R\rangle)$ for a self-avoiding homopolymer in a good solvent. In this case, the Redner- desCloizeaux [1, 2] distribution is given by,

$$P(r|\langle R\rangle) = A(r/\langle R\rangle)^{2+g}\exp(-B(r/\langle R\rangle)^{\delta}), \tag{1}$$

where $\langle R\rangle$ is the mean distance between two loci, and $g$ is the "correlation hole" exponent, and $\delta$ is related to the Flory exponent $\nu$ by $\delta = 1/(1-\nu)$. In good solvents, $\nu \approx 0.588$. The constants $A$ and $B$ can be calculated using the normalization condition, $\int \mathrm{d}r P(r|\langle R\rangle) = 1$. Given the value of $r_c$, the threshold distance for contact formation, the contact probability $P_c$ between the two loci is,

$$P_c = \int_0^{r_c} P(r|\langle R\rangle)\mathrm{d}r. \tag{2}$$

When the contact threshold is small compared to the size of the chain or the loop $r \ll \langle R\rangle$, the integral can be approximately evaluated using,

$$\begin{aligned} P_c &= \lim_{r_c\to 0}\int_0^{r_c} P(r)\mathrm{d}r \\ &= \lim_{r_c\to 0}\int_0^{r_c} A(r/\langle R\rangle)^{2+g}\exp(-B(r/\langle R\rangle)^{\delta})\mathrm{d}r, \\ &\sim \langle R\rangle^{-(3+g)}. \end{aligned} \tag{3}$$

Thus, the contact probability between two monomers, $P$, is related to the mean end-to-end distance $\langle R\rangle$ only through the scaling exponent $-(3+g)$. For ideal chain, $g = 0$, and thus we recover the asymptotically exact relation $P \sim \langle R \rangle^{-3}$. Note that $\langle R \rangle$ does depend on the genomic distance separating the two loci.

For a single polymer chain, there are three ways a contact between loci may be established [3]: i) the contact between two ends of the chain (Supplementary Figure 1a). ii) the contact between one end and a locus in the interior (Supplementary Figure 1b). iii) the contact between two loci in the interior of the chain (Supplementary Figure 1c). The correlation hole exponents corresponding to the three cases are $g_1 = 0.273$, $g_2 = 0.46$ and $g_3 = 0.71$ [3]. Thus, we have $P = \langle R \rangle^{-3.273}$, $P = \langle R \rangle^{-3.46}$ and $P = \langle R \rangle^{-3.71}$ for three cases. These rigorous values provide a bound for $g$, and should be viewed as a guide when considering the complicated case of chromosomes.

**Supplementary Note 2: Fitting the FISH data**

Here, we show how to fit the FISH data using Equations (3) in the main text. The integration of Supplementary Equation 1 gives the CDF$(R|\langle R \rangle)$,

$$\begin{aligned} \mathrm{CDF}(R|\langle R \rangle) &= \int_0^R P(r|\langle R \rangle) dr \\ &= 1 - \frac{A\langle R \rangle}{\delta} B^{-\frac{3+g}{\delta}} \Gamma\left(\frac{3+g}{\delta}, B\left(\frac{R}{\langle R \rangle}\right)^{\delta}\right), \end{aligned} \tag{4}$$

where $\Gamma(s, x)$ is the lower incomplete gamma function. Thus, the integral of Equation (3) in the main text with respect to $r$ may be written as,

$$\mathrm{CDF}(R) = \eta \mathrm{CDF}(R|\langle R_{1,mn} \rangle) + (1-\eta) \mathrm{CDF}(R|\langle R_{2,mn} \rangle). \tag{5}$$

Given the values of $g$, $\delta$ and $\langle R \rangle$, the two constants $A$ and $B$ are determined using the conditions: (1) The distribution (Supplementary Equation 1) is normalized, $\int_0^\infty \mathrm{d}r P(r|\langle R \rangle) = 1$. (2) The calculated value of the second moment $\langle R \rangle^2$, $\int_0^\infty \mathrm{d}r r P(r|\langle R \rangle) = \langle R \rangle$ should equal the measured value. With these two constraints, we obtain

$$\begin{aligned} A &= \frac{\delta}{\langle R \rangle} \frac{\Gamma^{3+g}((4+g)/\delta)}{\Gamma^{4+g}((3+g)/\delta)}, \\ B &= \frac{\Gamma^{\delta}((4+g)/\delta)}{\Gamma^{\delta}((3+g)/\delta)}, \end{aligned} \tag{6}$$

where $\Gamma(z)$ is the gamma function.

Using Supplementary Equation 6, Equation 4 can be further simplified as,

$$\mathrm{CDF}(R|\langle R\rangle) = 1 - \frac{\Gamma\big((3+g)/\delta, B(R/\langle R\rangle)^{\delta}\big)}{\Gamma((3+g)/\delta)}. \tag{7}$$

Using Supplementary Equations 5 and 7, we minimize the square of the difference between the calculated and the measured values of the CDFs. Using this procedure we calculated the values of $\eta$, $\langle R_{1,mn}\rangle$ and $\langle R_{2,mn}\rangle$ for all eight loci pairs for which FISH data were reported [4]. The best fit values of the three parameters are given in Supplementary Table 1. The goodness of the fits for different values of $g$ and $\delta$, corresponding to three different polymer models, are reported in Supplementary Table 2.

**Supplementary Note 3: Identification of the monomer type and loop anchors from experimental data**

We use the data provided in [4] to determine the location of the loop anchors. The locations of loops are provided in the file *GSE63525_GM12878_primary+ replicate_HiCCUPS_looplist_with_motifs.txt.gz* under the GEO accession number GSE63525. We only selected the loops with CTCF motifs "uniquely" called at both anchors (see Section VI.e.7 of the Extended Experimental Procedures of [4]). For each pair of CTCF loop anchors, we assign a harmonic constraint between the two corresponding loci.

**Supplementary Note 4: Fitting FISH data by assuming homogenous cell population**

To assess the quality of fits of the FISH data that assuming that the cell population is homogeneous, we use Supplementary Equation 7 with $g$ and $\delta$ as free parameters. Since for a homogenous population, the distribution of spatial distance can be normalized by dividing the spatial distance by the mean, which eliminates the parameter $\langle R\rangle$, leaving $g$ and $\delta$ as the only free parameters. Supplementary Figure 5 and Supplementary Table 3 show the result of the fits and the values of $g$ and $\delta$. First, the Kolmogorov-Smirnov statistics are poorer than for for fits obtained using two subpopulations. Second, the values the extracted values of $g$ and $\delta$ are unphysical. We, therefore, surmise that the FISH data cannot be reasonably fitted assuming the cell population is homogeneous.

**Supplementary Note 5 Kolmogorov-Smirnov test (K-S)**

The (K-S) test is a nonparametric test of the equality between two continuous probability distributions. Here, we use it to compare the experimental sample of the CDF with the theoretical probability distribution. The null hypothesis of the K-S test is that the data follow the reference distribution. The K-S test statistic is defined as,

$$D = \max_{1\le i\le N} \left| F(Y_i) - E(Y_i) \right| \tag{8}$$

where $F$ is the reference cumulative distribution and $E$ is the experimental (empirical) cumulative distribution. $Y_i$ are $N$ ordered (from smallest to largest) experimental measured data points. $E(Y_i) = n(i)/N$ where $n(i)$ is the number of data points less than $Y_i$. We use the *python* package *scipy* to compute the K-S statistics and its p-value.

**Supplementary Note 6: Fitting FISH data when heterogeneity is extensive**

In the results presented in Figure 4 in the main text, we assumed that chromosomes with or without CTCF loops may be categorized into two subpopulations, each with a characteristic mean distance (Fig.2 in the main text). Here, we generalize the theory using a continuous distribution of subpopulations, which is required in light of recent study [5]. Let us denote $P(\langle R\rangle)$ as the probability distribution of mean distance $\langle R\rangle$ characterizing a subpopulation. In Equation (3) in the main text, $P(\langle R\rangle)$ is assumed to be a linear combination of two Delta functions. This assumption, which is reasonable in the context of CTCF loops, may not hold in cases where cooperative interactions between loops are prominent or when there is extensive conformational heterogeneity. Here, we discuss how to analyze the FISH data without making any prior assumption about $P(\langle R\rangle)$. The generalization of Supplementary Equation 5 is,

$$\mathrm{CDF}(R) = \int_0^\infty \mathrm{d}\langle R\rangle P(\langle R\rangle)\mathrm{CDF}(R|\langle R\rangle). \tag{9}$$

If $P(\langle R\rangle) = \eta\delta(\langle R\rangle - \langle R_1\rangle) + (1-\eta)\delta(\langle R\rangle - \langle R_2\rangle)$, then we recover Supplementary Equation 5. However, extensive heterogeneity in chromosome organization implies that $\langle R\rangle$ could take arbitrary values with a distribution, $P(\langle R\rangle)$. The left side of Supplementary Equation 9 is the experimentally measured cumulative distribution function and $\mathrm{CDF}(R|\langle R\rangle)$ on the right hand side is given by Supplementary Equation 7. The goal is to solve for $P(\langle R\rangle)$ in Supplementary Equation 9, which is Fredholm integral equation of the first kind. In this work, we solve Supplementary Equation 9 using a discretization scheme on grid points $(R_j, \langle R\rangle_i)$. Supplementary Equation 9 is replaced by a summation approximately,

$$\mathrm{CDF}(R_j) = \sum_i \omega_i P(\langle R\rangle_i)\mathrm{CDF}(R_j|\langle R\rangle_i) \tag{10}$$

where $\omega_i$ are the weight coefficients for a quadrature formula. If we use small and equal grid size $\Delta_{\langle R\rangle} \to 0$, we can replace $\omega_i$ with $\Delta_{\langle R\rangle}$. Supplementary Equation 10 can be solved as a system of linear equations using non-negative Tikhonov regularization (see Supplementary Note 7).

**Supplementary Note 7: Non-negative Tikhonov regularization Method**

In this section, we show how to solve $P(\langle R\rangle_i)$ in Supplementary Equation 10 numerically. In order to simplify the notation, we denote $\mathrm{CDF}(R_j|\langle R\rangle_i) \equiv A$, $\mathrm{CDF}(R_j) \equiv b$ and $\Delta\langle R\rangle P(\langle R\rangle_i) \equiv x$. Then, Supplementary Equation 10 is written as,

$$Ax - b = 0 \tag{11}$$

Since Supplementary Equation 9 is an integral equation for which solutions may not be unique. For such an ill-posed problem, Supplementary Equation 10 (or Supplementary Equation 11) is usually solved using the Tikhonov regularization method, which is to solve,

$$\min(||Ax - b||_2^2 + \alpha^2||x||_2^2), \tag{12}$$

where $\alpha$ is a tuned parameter controlling the smoothness of solution $x$. For our problem, we also need an additional non-negative constraint on $x$ since $x$ is a probability density function. Thus, we want to solve Supplementary Equation 12 subject to $x \geq 0$. Let us construct the matrix,

$$C = \begin{bmatrix} A \\ \alpha I \end{bmatrix} \tag{13}$$

and the vector,

$$d = \begin{bmatrix} b \\ 0 \end{bmatrix} \tag{14}$$

where $I$ is the identity matrix. Solving Supplementary Equation 13 subject to $x \geq 0$ is equivalent to solving,

$$\min||Cx - d||_2^2, \text{ subject to } x \geq 0. \tag{15}$$

The above equation is a non-negative least square (NNLS) problem, which can be solved using an active set algorithm [6]. To solve the Supplementary Equation 15, the value of $\alpha$, which is a controlled parameter, needs to be provided. From a graphical perspective, $\alpha$ controls the smoothness of the solution, with $x$ being smoother for larger $\alpha$. The statistical significance of $\alpha$ lies in its ability to control the trade-off between the goodness of fit and the extent of over-fitting. To choose the value of $\alpha$ in a systematic way, we follow the procedure demonstrated in [7]. The goodness of fits is measured by the residue norm $||Ax - b||_2^2$ and the solution norm $||x||_2^2$ is used as a proxy to the extent of over-fitting. The L-curve is the function between $||Ax - b||_2^2$ and $||x||_2^2$ for different values of $\alpha$. The optimal value of $\alpha$ is located where the L-curve has largest curvature. Supplementary Figure 7 shows an example of the procedure. We solve Supplementary Equation 15 using the optimal value of $\alpha$. In practice, we use the function provided in PYTHON *scipy* package to solve Supplementary Equation 15.

**Supplementary Note 8: Correlation between two loops**

In this section, we derive the conditional probability of contact between $m$ and $n$ loci given that $k$ and $l$ loci are in contact for GRMC. Let us denote the distances between $m$ and $n$, $k$ and $l$ as $R_1$ and $R_2$, respectively. The goal is to derive $\text{Prob}(R_1 \leq r_c | R_2 \leq r_c)$. From Bayes' theorem, we know that

$$\text{Prob}(R_1 \leq r_c | R_2 \leq r_c) = \frac{\text{Prob}(R_1 \leq r_c, R_2 \leq r_c)}{\text{Prob}(R_2 \leq r_c)} \tag{16}$$

In order to calculate $\text{Prob}(R_1 \leq r_c, R_2 \leq r_c)$ in the above equation, recall that

$$\begin{aligned} \mathbf{R}_1 &= \sum_{p=0}^{N-1} (V_{pm} - V_{pn}) \mathbf{X}_p, \\ \mathbf{R}_2 &= \sum_{p=0}^{N-1} (V_{pk} - V_{pl}) \mathbf{X}_p, \end{aligned} \tag{17}$$

where $\mathbf{X}_p$ are the normal modes, and $V_{pm}$,$V_{pn}$,$V_{pk}$ and $V_{pl}$ are the elements in the orthonormal matrix (see Equation (7) in the main text). We can write Supplementary Equation 17 given above as,

$$\underbrace{\begin{pmatrix} R_{1,\alpha} \\ R_{2,\alpha} \end{pmatrix}}_{\mathbf{R}_\alpha} = \underbrace{\begin{pmatrix} V_{0,m} - V_{0,n} & V_{1,m} - V_{1,n} & \cdots & V_{N-1,m} - V_{N-1,n} \\ V_{0,k} - V_{0,l} & V_{1,k} - V_{1,l} & \cdots & V_{N-1,k} - V_{N-1,l} \end{pmatrix}}_{\mathbf{V}_{mn,kl}} \underbrace{\begin{pmatrix} X_{0,\alpha} \\ X_{1,\alpha} \\ \vdots \\ X_{N-1,\alpha} \end{pmatrix}}_{\mathbf{X}_\alpha} \tag{18}$$

where $\alpha = x, y, z$. The normal modes $\mathbf{X}_\alpha$ have multivariate normal distribution, $\mathbf{X}_\alpha \sim \mathcal{N}(\mathbf{0}, \mathbf{\Sigma})$ where $\mathbf{0} = (0, 0, \cdots, 0)^T$, and $\mathbf{\Sigma} = \mathrm{diag}(-\frac{k_BT}{\lambda_0}, -\frac{k_BT}{\lambda_1}, \cdots, -\frac{k_BT}{\lambda_{N-1}})$. Hence, $\mathbf{R}_\alpha$ also obey the multivariate normal distribution, $\mathbf{R}_\alpha \sim \mathcal{N}(\mathbf{0}, \mathbf{V}_{mn,kl}\mathbf{\Sigma}\mathbf{V}_{mn,kl}^T) \equiv \mathcal{N}(\mathbf{0}, \mathbf{M})$. We have,

$$\mathbf{M} = \mathbf{V}_{mn,kl}\mathbf{\Sigma}\mathbf{V}_{mn,kl}^T = \begin{pmatrix} M_{11} & M_{12} \\ M_{21} & M_{22} \end{pmatrix} \tag{19}$$

where $M_{11} = -\sum_{p=0}^{N-1}(V_{pm} - V_{pn})^2 \frac{k_BT}{\lambda_p} \equiv \sigma_1^2$, $M_{22} = -\sum_{p=0}^{N-1}(V_{pk} - V_{pl})^2 \frac{k_BT}{\lambda_p} \equiv \sigma_2^2$ and $M_{12} = M_{21} = -\sum_{p=0}^{N-1}(V_{pm} - V_{pn})(V_{pk} - V_{pl})\frac{k_BT}{\lambda_p}$. Note that $\mathbf{M}$ is a covariance matrix. Thus, we denote $M_{12} = M_{21} = \rho\sigma_1\sigma_2$ where $\rho$ is the correlation coefficient between $R_{1,\alpha}$ and $R_{2,\alpha}$.

So far, we have established that $\mathbf{R}_\alpha$ has a bivariate normal distribution with mean zero and covariance matrix $\mathbf{M}$. Note that $R_1 = \sqrt{\sum_{\alpha=x,y,z} R_{1,\alpha}^2}$ and $R_2 = \sqrt{\sum_{\alpha=x,y,z} R_{2,\alpha}^2}$. We want to know the joint probability distribution of $R_1$ and $R_2$, $p(R_1, R_2)$. This distribution is bivariate central chi distribution, also referred to as generalized Rayleigh distribution [8]. It has a closed form [8]

$$p(R_1, R_2) = A(R_1R_2)^{3/2} e^{-S/2} I_{1/2}\left(\frac{|\rho| R_1 R_2}{\sigma_1\sigma_2(1-\rho^2)}\right) \tag{20}$$

where $A = \frac{\sqrt{2/\pi}}{(\sigma_1\sigma_2)^{5/2}(1-\rho^2)|\rho|^{1/2}}$, $S = \frac{R_1^2}{\sigma_1^2(1-\rho^2)} + \frac{R_2^2}{\sigma_2^2(1-\rho^2)}$ and $I_{1/2}(\cdot)$ is the modified Bessel function of the first kind. Thus, $\mathrm{Prob}(R_1 \le r_c, R_2 \le r_c)$ in Supplementary Equation 16 can be readily computed as,

$$\mathrm{Prob}(R_1 \le r_c, R_2 \le r_c) = \int_0^{r_c}\int_0^{r_c} \mathrm{d}R_1 \mathrm{d}R_2\, p(R_1, R_2) \tag{21}$$

There is no simple form for the integral of Supplementary Equation 16. However it can be computed for small $r_c$ limit by using the fact that $\lim_{z\to 0} I_{1/2}(z) = \frac{(\frac{1}{2}z)^{1/2}}{\Gamma(3/2)}$.

$$\begin{aligned}
&\lim_{r_c\to 0}\int_0^{r_c}\int_0^{r_c} \mathrm{d}R_1 \mathrm{d}R_2 p(R_1,R_2) \\
&= A\sqrt{\frac{2}{\pi}}\left(\frac{|\rho|}{\sigma_1\sigma_2(1-\rho^2)}\right)^{1/2}\int_0^{r_c} \mathrm{d}R_1^2 e^{-\frac{R_1^2}{2\sigma_1^2(1-\rho^2)}}\int_0^{r_c} \mathrm{d}R_2^2 e^{-\frac{R_2^2}{2\sigma_2^2(1-\rho^2)}} \\
&\approx (1-\rho^2)^{3/2}\left(\sqrt{\frac{2}{\pi}}\frac{1}{3}\frac{(r_c/\sqrt{1-\rho^2})^3}{\sigma_1^3}\right)\left(\sqrt{\frac{2}{\pi}}\frac{1}{3}\frac{(r_c/\sqrt{1-\rho^2})^3}{\sigma_2^3}\right)
\end{aligned} \tag{22}$$

For small $r_c$, we have $\mathrm{Prob}(R_2 \le r_c) = \sqrt{\frac{2}{\pi}}\frac{1}{3}\frac{r_c^3}{\sigma_2^3}$. Finally, we have

$$\mathrm{Prob}(R_1 \le r_c | R_2 \le r_c) = \frac{\mathrm{Prob}(R_1 \le r_c, R_2 \le r_c)}{\mathrm{Prob}(R_2 \le r_c)} = \sqrt{\frac{2}{\pi}}\frac{1}{3}\frac{(r_c/\sqrt{1-\rho^2})^3}{\sigma_1^3} \tag{23}$$

Recall that $\langle R_1\rangle = 2\sqrt{\frac{2}{\pi}}\sigma_1$. Thus, $\mathrm{Prob}(R_1 \le r_c | R_2 \le r_c) \approx \frac{(r_c/\sqrt{1-\rho^2})^3}{\langle R_1\rangle^3}$. Given that $\mathrm{Prob}(R_1 \le r_c) \approx \frac{r_c^3}{\langle R_1\rangle^3}$, the conditional probability differs from the marginal probability only by a factor $1/(1-\rho^2)^{3/2}$. This result is intuitive. When $\rho = 0$, there is no correlation between $R_1$ and $R_2$, the conditional probability simply is the marginal probability. When $\rho \neq 0$, pinning the contact between $k$ and $l$ always increases the contact probability between $m$ and $n$, essentially by proximity rule considered in an entirely different context [9]. The scaling exponent between the conditional contact probability and mean distance remains the same ($= 3$).

| | $\eta$ | $\langle R_{1,mn}\rangle(\mu m)$ | $\langle R_{2,mn}\rangle(\mu m)$ | Kolmogorov-Smirnov statistics | p-value |
|---|---|---|---|---|---|
| peak1-control | 0.55 | 0.63 | 1.06 | 0.0350 | 0.999 |
| peak1-loop | 0.36 | 0.24 | 0.56 | 0.0619 | 0.834 |
| peak2-control | 0.63 | 0.47 | 1.04 | 0.0522 | 0.977 |
| peak2-loop | 0.75 | 0.33 | 1.07 | 0.110 | 0.248 |
| peak3-control | 0.97 | 0.67 | 4.08 | 0.0561 | 0.902 |
| peak3-loop | 0.91 | 0.35 | 1.64 | 0.0507 | 0.954 |
| peak4-control | 0.27 | 0.48 | 1.25 | 0.0633 | 0.988 |
| peak4-loop | 0.42 | 0.30 | 1.21 | 0.0657 | 0.982 |

Supplementary Table I: Values of the optimal parameters obtained by fitting the FISH data (plotted in Supplementary Figure 2 (red curves)) to Equation (3) using our theory. The parameters $\eta$, $\langle R_{1,mn}\rangle(\mu m)$, and $\langle R_{2,mn}\rangle(\mu m)$ are defined in the main text. It is interesting that the values of $\langle R_{1,mn}\rangle$ for the peak-loop positions are similar ($\approx 0.3\mu$m). Kolmogorov-Smirnov statistics and their p-values are also reported.

| g | $\delta$ | RE |
|---|---|---|
| 1 | 5/4 | 0.00392 |
| 0 | 2 | 0.00397 |
| 0.71 | 5/2 | 0.00762 |

Supplementary Table II: Residual error (RE) for fits of theory to the FISH data using three different sets of $g$ and $\delta$. We define RE as RE $= \sum_j (1/N_j) \sum_i^{N_j} (y_i - f(x_i))^2$ where $y_i$ is the $i^{th}$ value of the measured data, $f(x_i)$ is the fit value for $y_i$. The sum is over all the data points from all the eight curves marked by index $j$ ($j = 1, 2, \cdots, 8$). $N_j$ is the number of data points of $j^{th}$ curve. The values of $g$ and $\delta$ in the first row corresponds to chromosomes, while those in the second and third rows are for the Rouse model and a polymer in a good solvent, respectively. The smallest error is for the exponents describing the chromosome model. Surprisingly, the RE value for the unphysical Rouse model is also low.

| | g | $\delta$ | Kolmogorov-Smirnov statistics | p-value |
|---|---|---|---|---|
| peak1-control | 8.68 | 0.30 | 0.0346 | 0.999 |
| peak1-loop | 115.37 | 0.017 | 0.0716 | 0.678 |
| peak2-control | 111.38 | 0.017 | 0.0923 | 0.458 |
| peak2-loop | 108.70 | 0.012 | 0.118 | 0.180 |
| peak3-control | 204.82 | 0.015 | 0.113 | 0.132 |
| peak3-loop | 180.84 | 0.010 | 0.148 | 0.019 |
| peak4-control | -0.99 | 1.42 | 0.0558 | 0.997 |
| peak4-loop | -1.76 | 1.26 | 0.0863 | 0.850 |

Supplementary Table III: Values of the optimal $g$ and $\delta$ obtained by fitting the FISH data (Supplementary Figure 5) assuming that the cell population is homogenous (Supplementary Equation 7). The best fit values of $g$ and $\delta$ are unphysical. The results of the Kolmogorov-Smirnov test are are inferior to the results reported in Table S1.

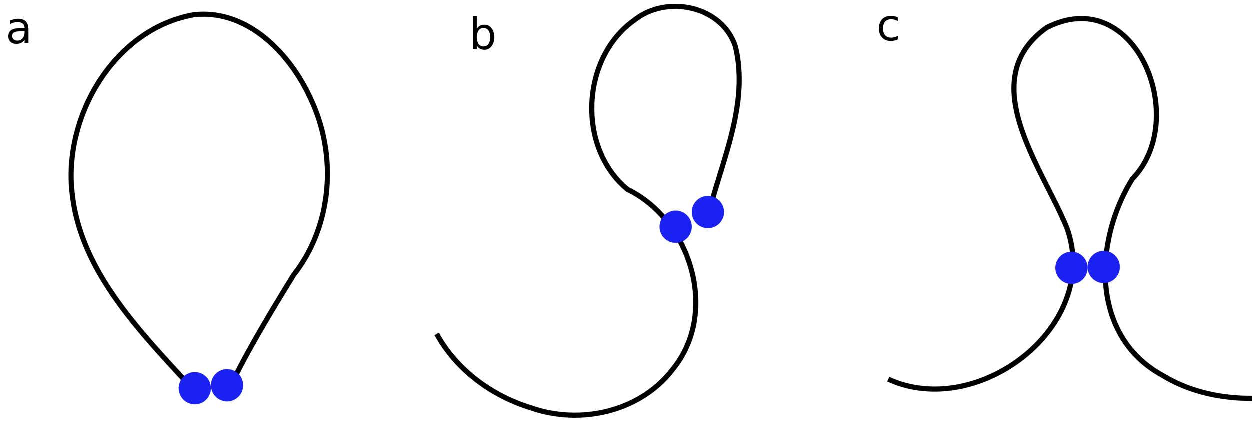


Supplementary Figure 1: Three possibilities for contact formation between two loci in a polymer. **(a)** Contact formation between the two ends. **(b)** Contact formation between one end and a locus in the interior. **(c)** Contact formation between two loci located in the interior of a polymer. Although the relation between $P$ and $\langle R \rangle$ decreases as power law the value of the exponent is different in the three scenarios (see the Supplementary Note 1 for the precise values of a polymer in good solvent).

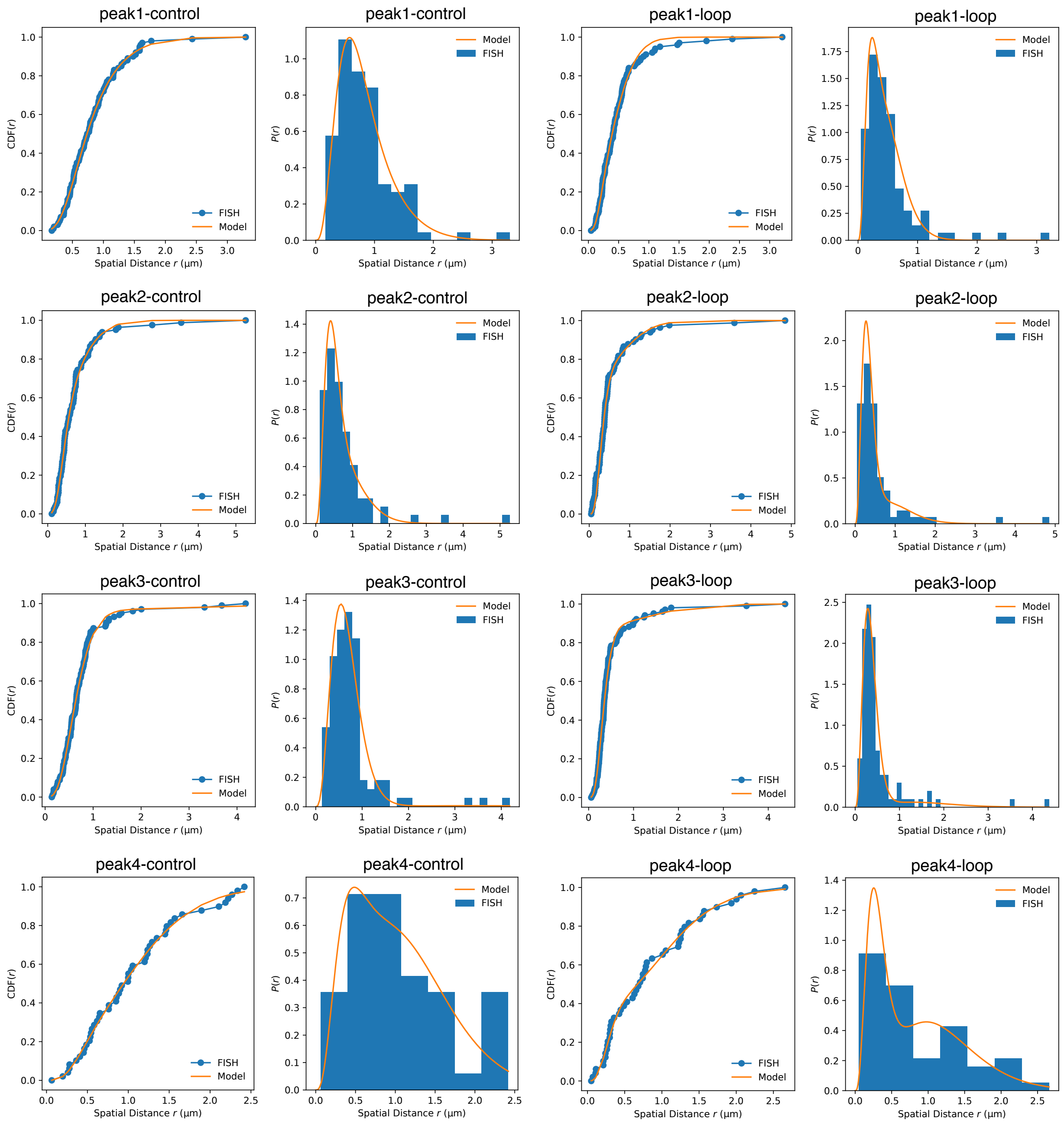


Supplementary Figure 2: Fit of the CDF($R$) (using Supplementary Equation 5 with values of $g = 1$ and $\delta = 5/4$) to the experimental data (blue dots) [4]. Orange lines are the fits. The parameters obtained from the fits for the eight loci pair are summarized in Supplementary Table 1. The probability density distribution (PDF) obtained using the fit parameters are also plotted along with experimental PDF. The excellent agreement between theory and experiments is self-evident.

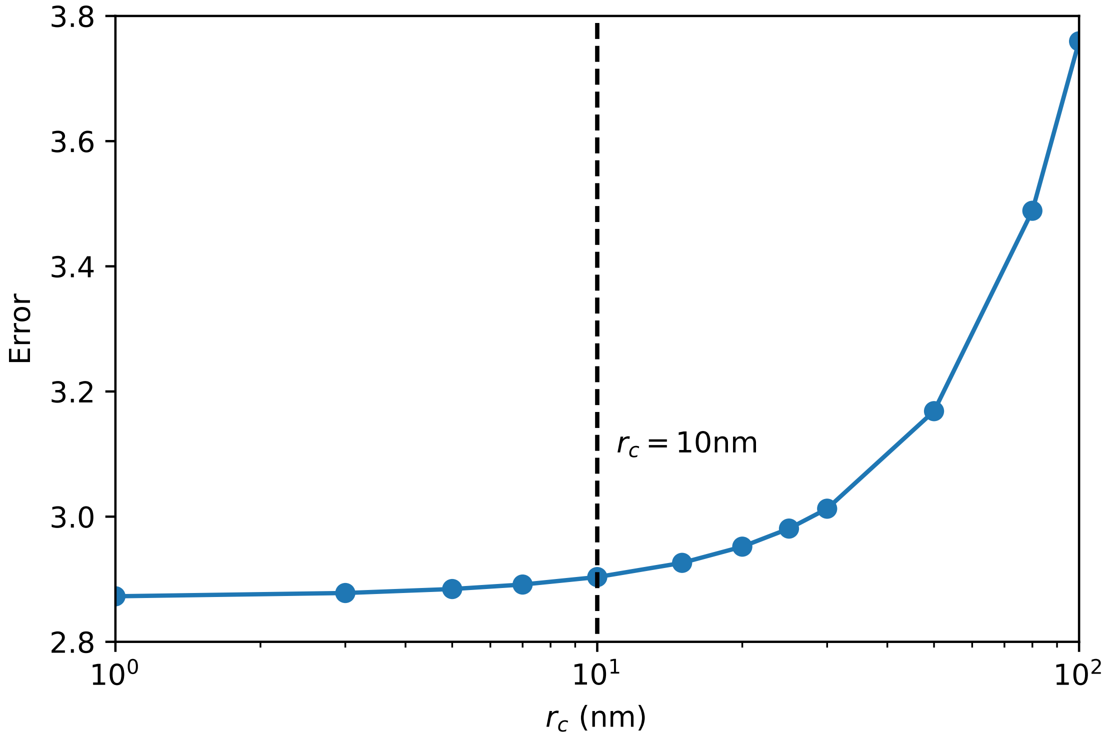


Supplementary Figure 3: Predicted error in the contact probability with respect to Hi-C measurements as function of $r_c$. The error is computed using $\sum_i \Big[ \log(P_i^{\mathrm{Model}}/\langle P^{\mathrm{Model}}\rangle) - \log(P_i^{\mathrm{HiC}}/\langle P^{\mathrm{HiC}}\rangle) \Big]^2$. We use logarithm for the errors because the contact probability for different pairs can differ by orders of magnitude (see Fig.3 and 4(b) in the main text for example).

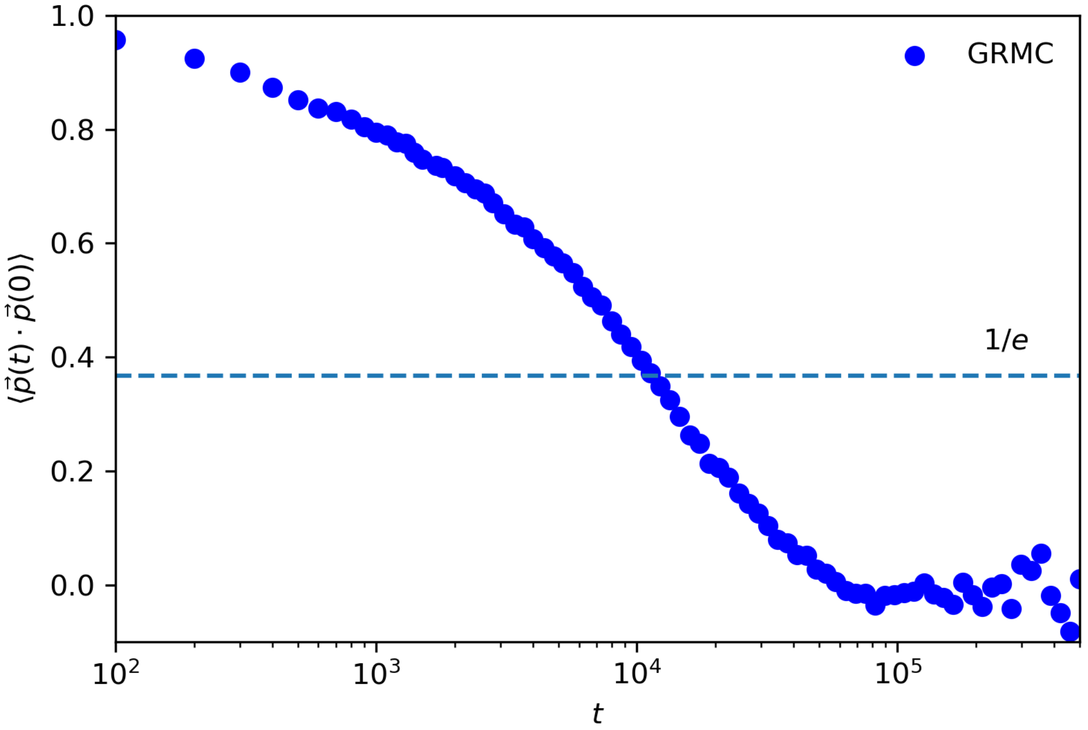


Supplementary Figure 4: Autocorrelation of the end-to-end vector. The end-to-end vector at time $t$ is denoted as $\vec{p}(t)$. The relaxation time $\tau$ (when $\langle \vec{p}(t) \cdot \vec{p}(0) \rangle = 1/e$) for GRMC is about $10^6$ time steps. The total duration of one trajectory in the production runs is $10^8$ timesteps and we simulated a total number of 10 independent trajectories. Thus, our simulation produces about 800 independent snapshots for GRMC. This figure is meant illustrate that both the length and number of trajectories are adequate in obtaining converged results in the simulations.

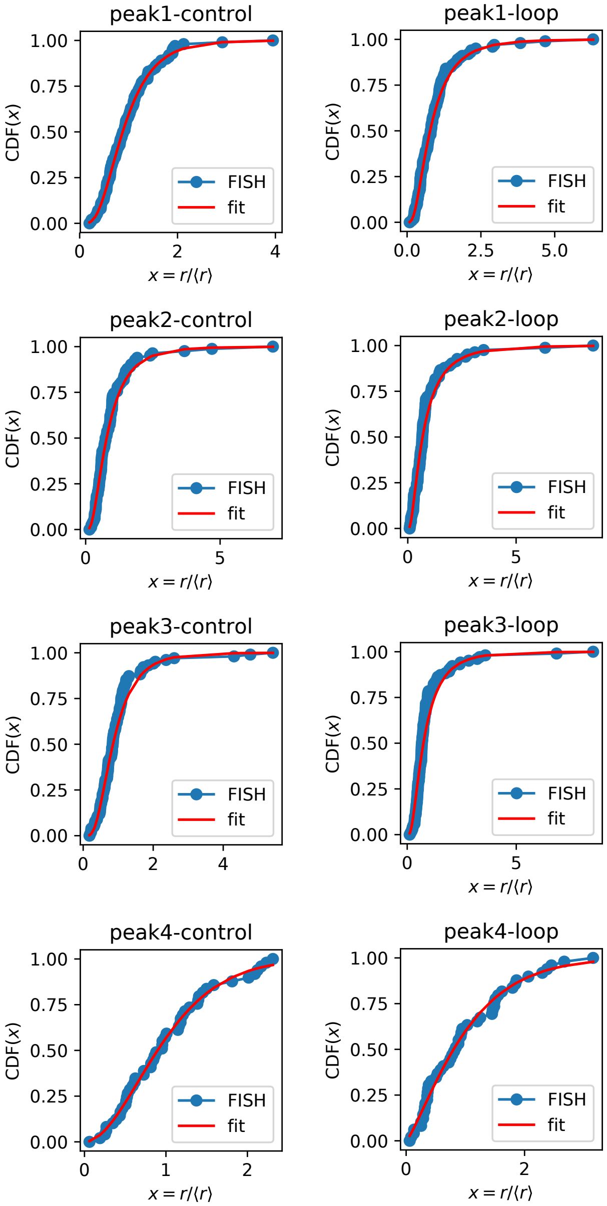


Supplementary Figure 5: Fit of the CDF$(R|\langle R\rangle)$ using Supplementary Equation 7 to the experimental data (blue dots) [4]. Orange lines are the fits. The values of $g$ and $\delta$ obtained from the fits for the eight loci pair are summarized in the Supplementary Table 3. Although the fits assuming homogeneous cell population are good, the optimal values of $g$ and $\delta$ are unphysical (see Supplementary Table 3).

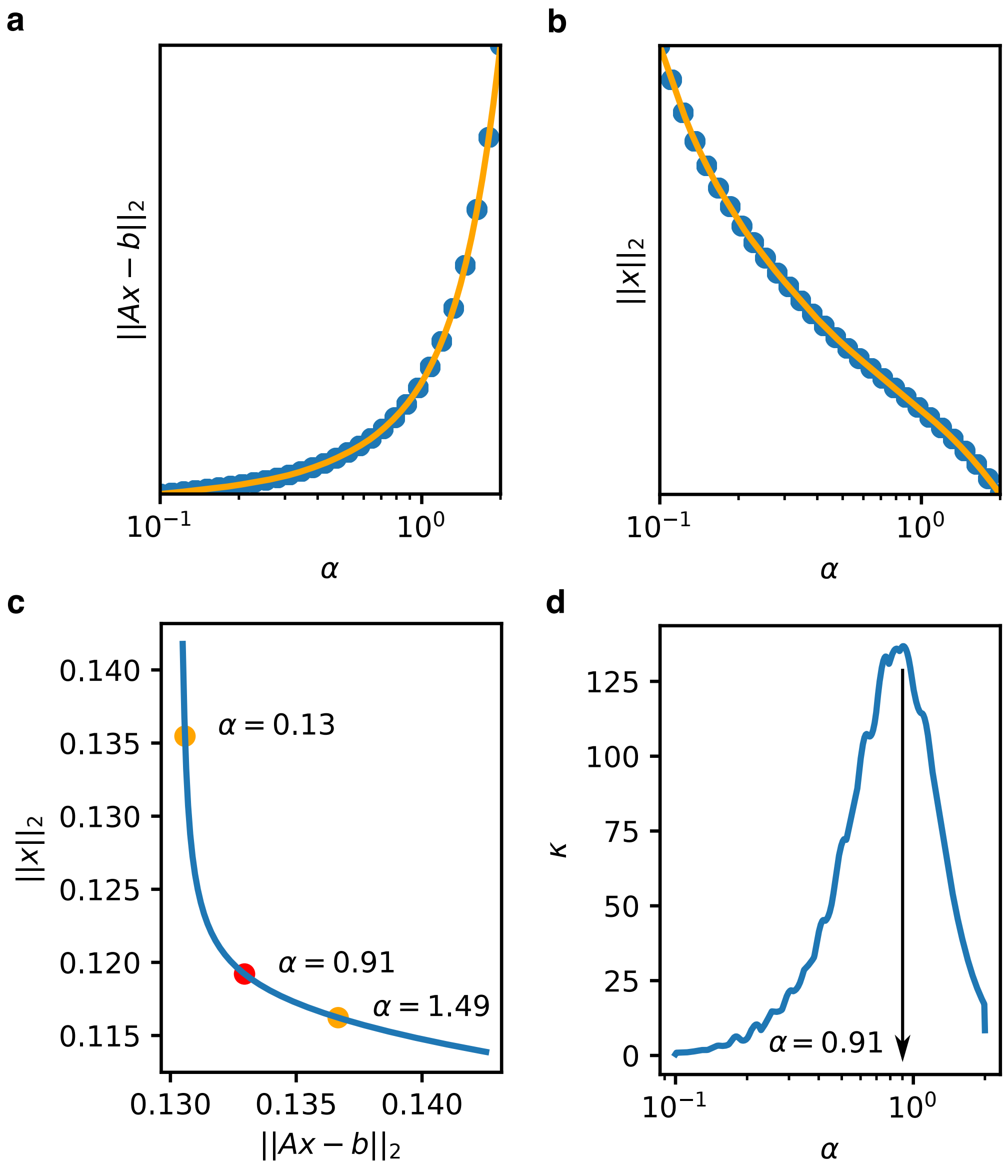


Supplementary Figure 6: An example of the procedure for choosing the value of parameter $\alpha$. **(a)** Residue norm $||Ax - b||_2$ as a function of $\alpha$. **(b)** Solution norm $||x||_2$ as a function of $\alpha$. **(c)** The L-curve, plotted as the function between the residue norm $||Ax - b||_2$ and the solution norm $||x||_2$. Three points on L-curve are marked, each one indicated by its corresponding value of $\alpha$. **(d)** The curvature $\kappa$ of L-curve is calculated as a function of $\alpha$. The value of $\alpha$ is chosen to be the one at which the curvature $\kappa$ is maximized.

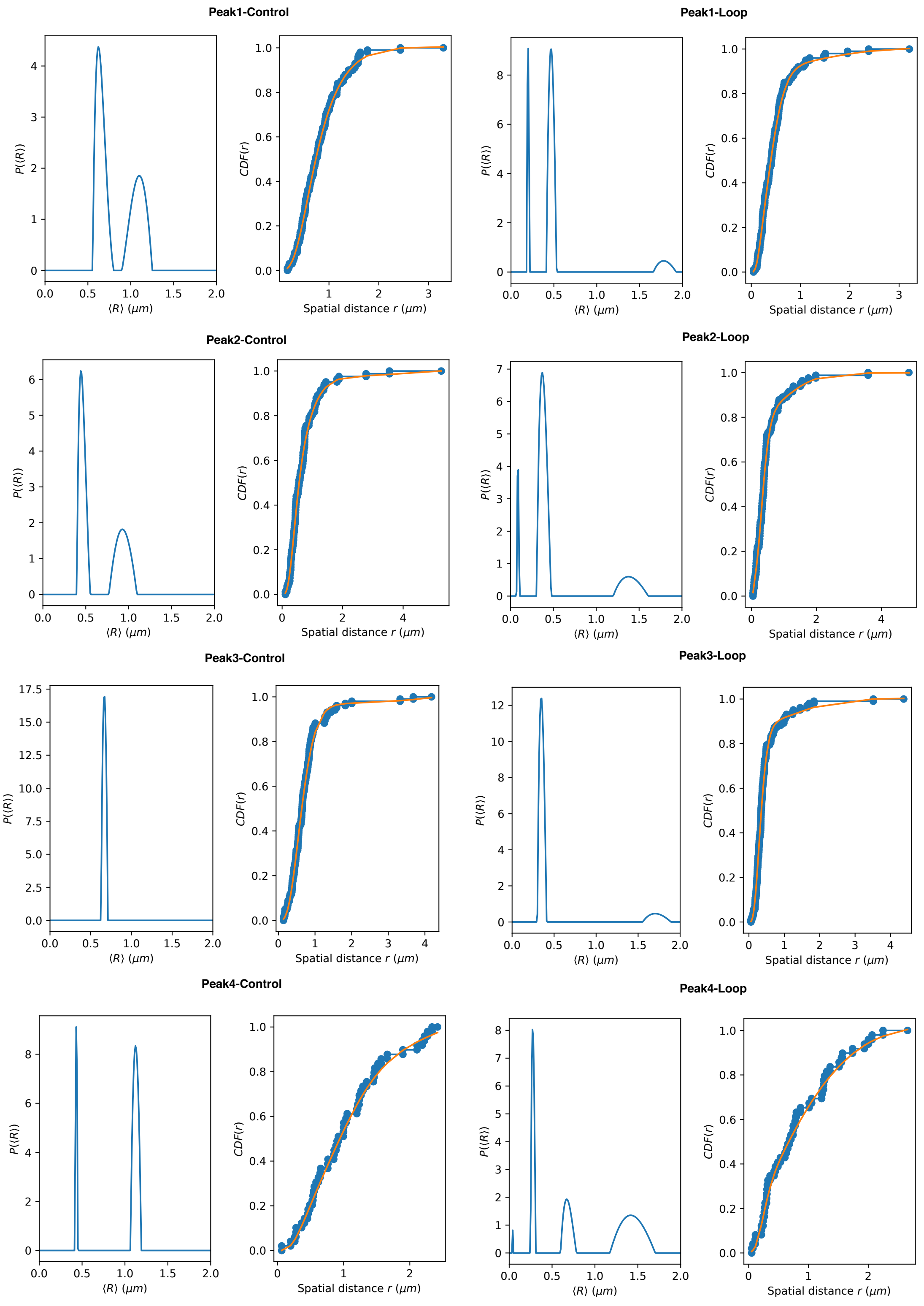
Peak1-Control
Peak1-Loop
Peak2-Control
Peak2-Loop
Peak3-Control
Peak3-Loop
Peak4-Control
Peak4-Loop
$P(\langle R \rangle)$
$\langle R \rangle$ ($\mu m$)
$CDF(r)$
Spatial distance $r$ ($\mu m$)

Supplementary Figure 7: Fit of CDF$(r)$ using Supplementary Equation 9 to the experimental data (blue dots) [4]. Orange lines are the calculated results using our theory. From the fit, the distribution $P(\langle R \rangle)$ given in the integral equation (Supplementary Equation 9) is inversely solved using the non-negative Tikhonov Regularization method. As shown here, multi-peaks feature naturally emerge without assuming the functional form of $P(\langle R \rangle)$ as in Supplementary Equation 5. We set $g = 1$ and $\delta = 5/4$.

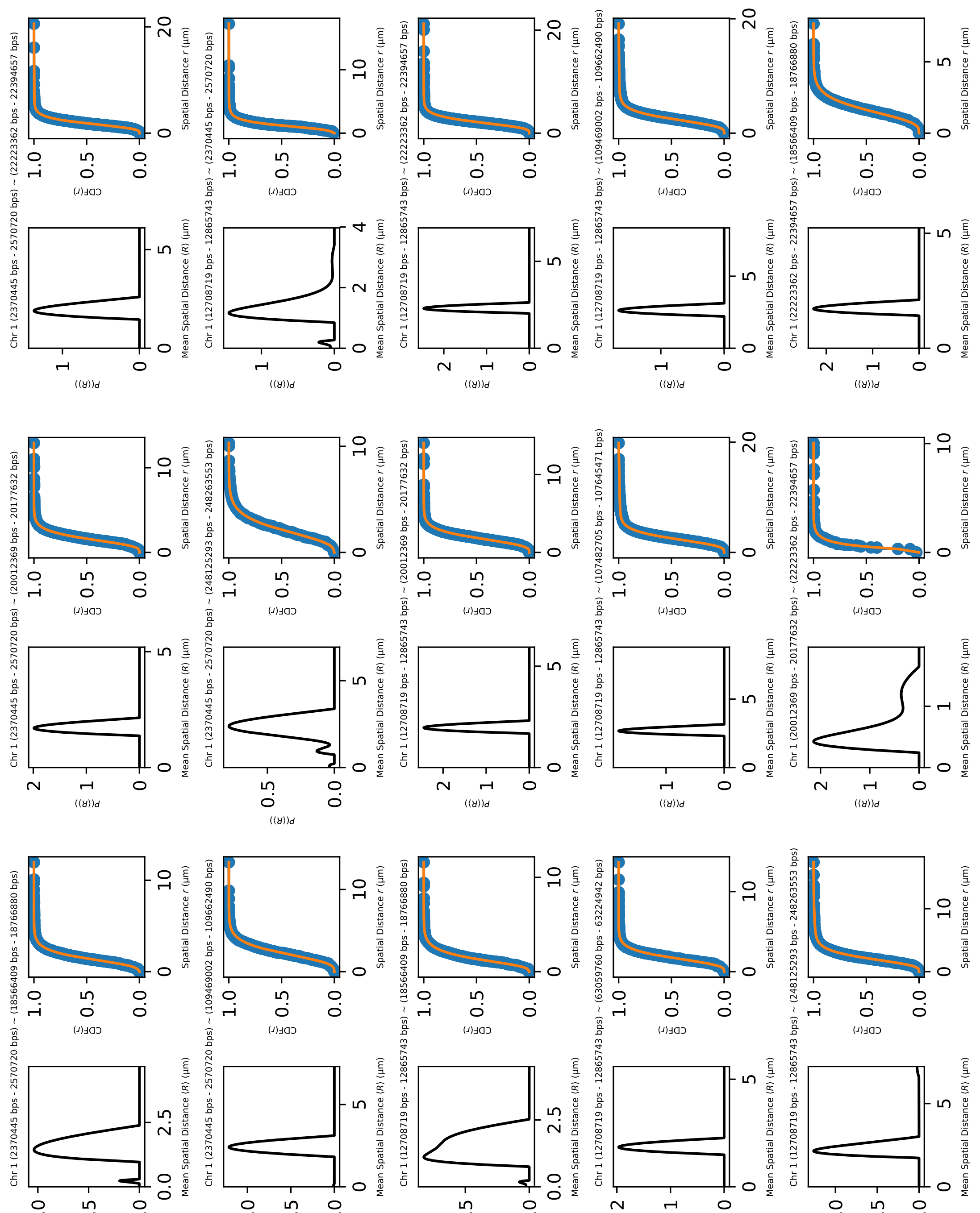
a
Chr 1 (2370445 bps - 2570720 bps) ~ (18566409 bps - 18766880 bps)
Chr 1 (2370445 bps - 2570720 bps) ~ (20012369 bps - 20177632 bps)
Chr 1 (2370445 bps - 2570720 bps) ~ (22223362 bps - 22394657 bps)
Chr 1 (2370445 bps - 2570720 bps) ~ (109469002 bps - 109662490 bps)
Chr 1 (2370445 bps - 2570720 bps) ~ (248125293 bps - 248263553 bps)
Chr 1 (12708719 bps - 12865743 bps) ~ (2370445 bps - 2570720 bps)
Chr 1 (12708719 bps - 12865743 bps) ~ (18566409 bps - 18766880 bps)
Chr 1 (12708719 bps - 12865743 bps) ~ (20012369 bps - 20177632 bps)
Chr 1 (12708719 bps - 12865743 bps) ~ (22223362 bps - 22394657 bps)
Chr 1 (12708719 bps - 12865743 bps) ~ (63059760 bps - 63224942 bps)
Chr 1 (12708719 bps - 12865743 bps) ~ (107482705 bps - 107645471 bps)
Chr 1 (12708719 bps - 12865743 bps) ~ (109469002 bps - 109662490 bps)
Chr 1 (12708719 bps - 12865743 bps) ~ (248125293 bps - 248263553 bps)
Chr 1 (20012369 bps - 20177632 bps) ~ (22223362 bps - 22394657 bps)
Chr 1 (22223362 bps - 22394657 bps) ~ (18566409 bps - 18766880 bps)
P(⟨R⟩)
Mean Spatial Distance ⟨R⟩ (μm)
CDF(r)
Spatial Distance r (μm)

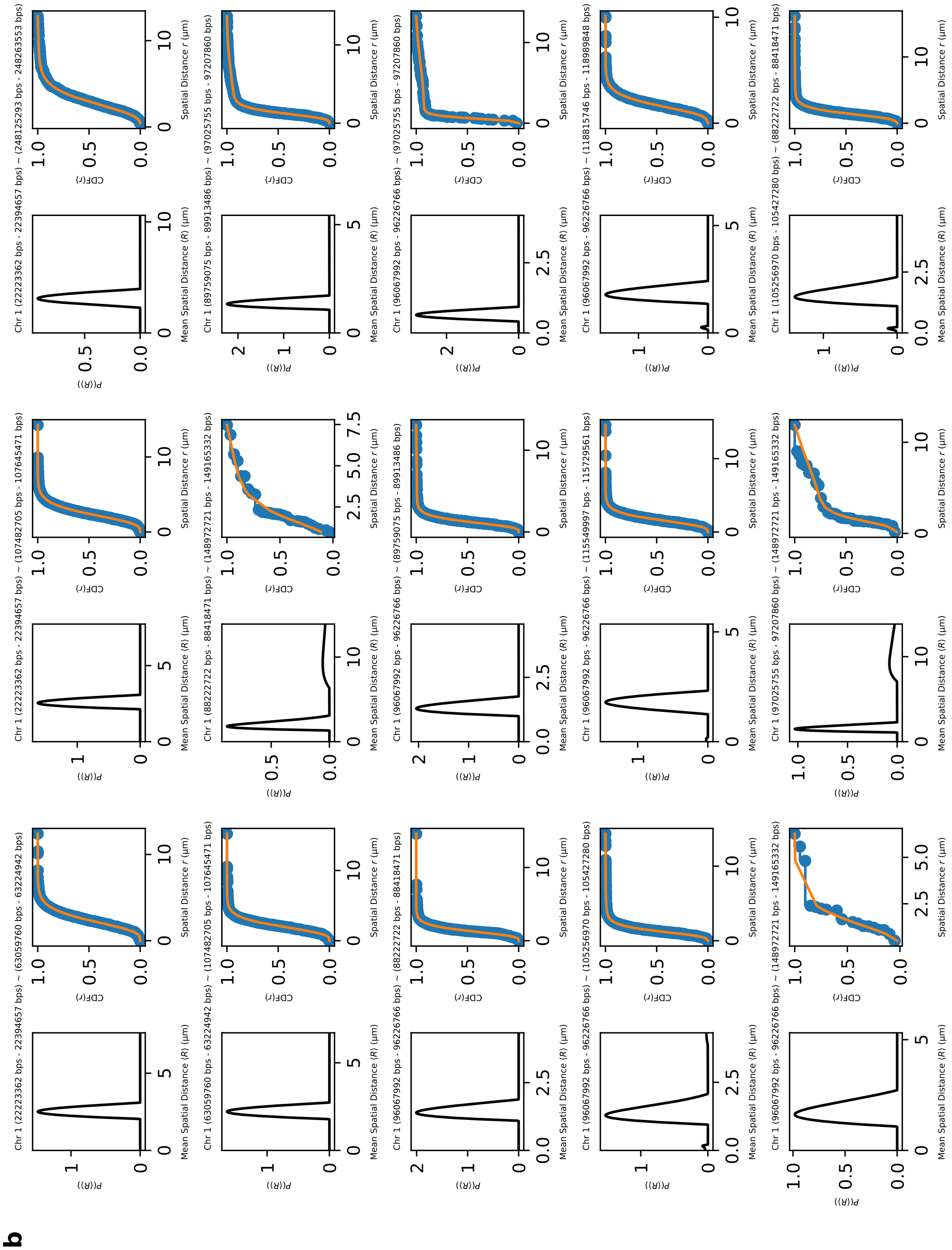
b

**c**

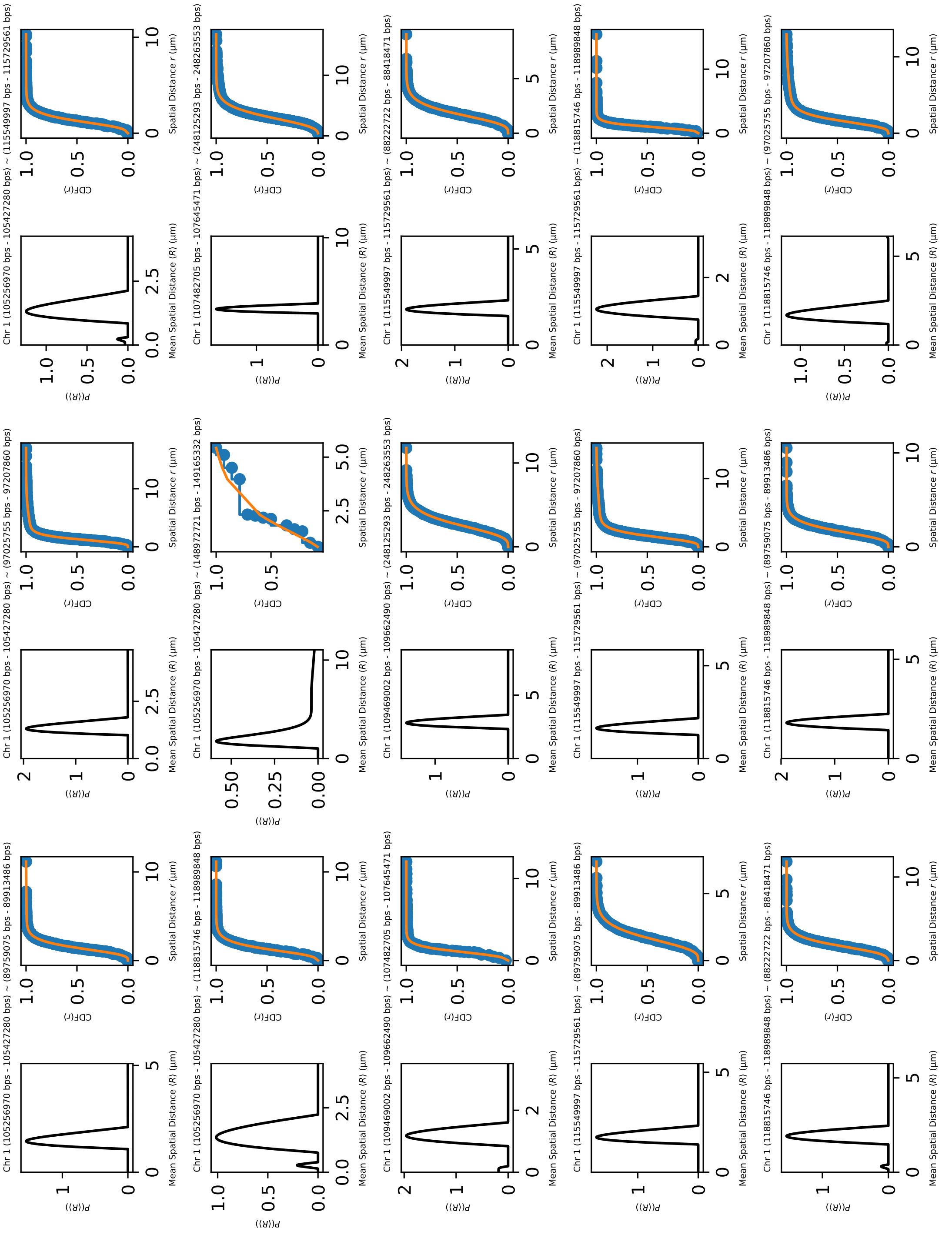
Chr 1 (105256970 bps - 105427280 bps) ~ (89759075 bps - 89913486 bps)
Chr 1 (105256970 bps - 105427280 bps) ~ (97025755 bps - 97207860 bps)
Chr 1 (105256970 bps - 105427280 bps) ~ (115549997 bps - 115729561 bps)
Chr 1 (105256970 bps - 105427280 bps) ~ (118815746 bps - 118989848 bps)
Chr 1 (105256970 bps - 105427280 bps) ~ (148972721 bps - 149165332 bps)
Chr 1 (107482705 bps - 107645471 bps) ~ (248125293 bps - 248263553 bps)
Chr 1 (109469002 bps - 109662490 bps) ~ (107482705 bps - 107645471 bps)
Chr 1 (109469002 bps - 109662490 bps) ~ (248125293 bps - 248263553 bps)
Chr 1 (115549997 bps - 115729561 bps) ~ (88222722 bps - 88418471 bps)
Chr 1 (115549997 bps - 115729561 bps) ~ (89759075 bps - 89913486 bps)
Chr 1 (115549997 bps - 115729561 bps) ~ (97025755 bps - 97207860 bps)
Chr 1 (115549997 bps - 115729561 bps) ~ (118815746 bps - 118989848 bps)
Chr 1 (118815746 bps - 118989848 bps) ~ (88222722 bps - 88418471 bps)
Chr 1 (118815746 bps - 118989848 bps) ~ (89759075 bps - 89913486 bps)
Chr 1 (118815746 bps - 118989848 bps) ~ (97025755 bps - 97207860 bps)
P(⟨R⟩)
Mean Spatial Distance ⟨R⟩ (μm)
CDF(r)
Spatial Distance r (μm)

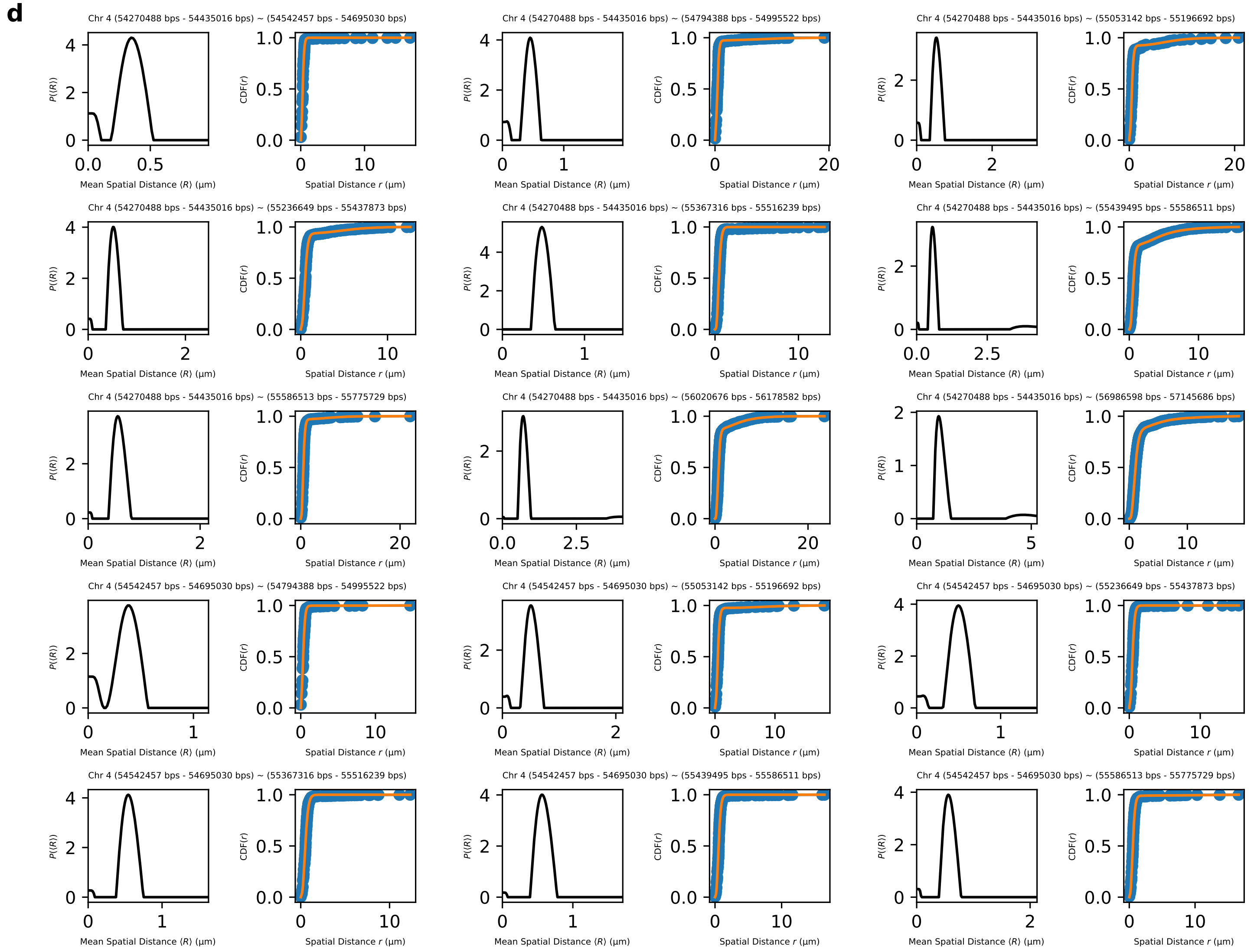
d
Chr 4 (54270488 bps - 54435016 bps) ~ (54542457 bps - 54695030 bps)
Chr 4 (54270488 bps - 54435016 bps) ~ (54794388 bps - 54995522 bps)
Chr 4 (54270488 bps - 54435016 bps) ~ (55053142 bps - 55196692 bps)
Chr 4 (54270488 bps - 54435016 bps) ~ (55236649 bps - 55437873 bps)
Chr 4 (54270488 bps - 54435016 bps) ~ (55367316 bps - 55516239 bps)
Chr 4 (54270488 bps - 54435016 bps) ~ (55439495 bps - 55586511 bps)
Chr 4 (54270488 bps - 54435016 bps) ~ (55586513 bps - 55775729 bps)
Chr 4 (54270488 bps - 54435016 bps) ~ (56020676 bps - 56178582 bps)
Chr 4 (54270488 bps - 54435016 bps) ~ (56986598 bps - 57145686 bps)
Chr 4 (54542457 bps - 54695030 bps) ~ (54794388 bps - 54995522 bps)
Chr 4 (54542457 bps - 54695030 bps) ~ (55053142 bps - 55196692 bps)
Chr 4 (54542457 bps - 54695030 bps) ~ (55236649 bps - 55437873 bps)
Chr 4 (54542457 bps - 54695030 bps) ~ (55367316 bps - 55516239 bps)
Chr 4 (54542457 bps - 54695030 bps) ~ (55439495 bps - 55586511 bps)
Chr 4 (54542457 bps - 54695030 bps) ~ (55586513 bps - 55775729 bps)

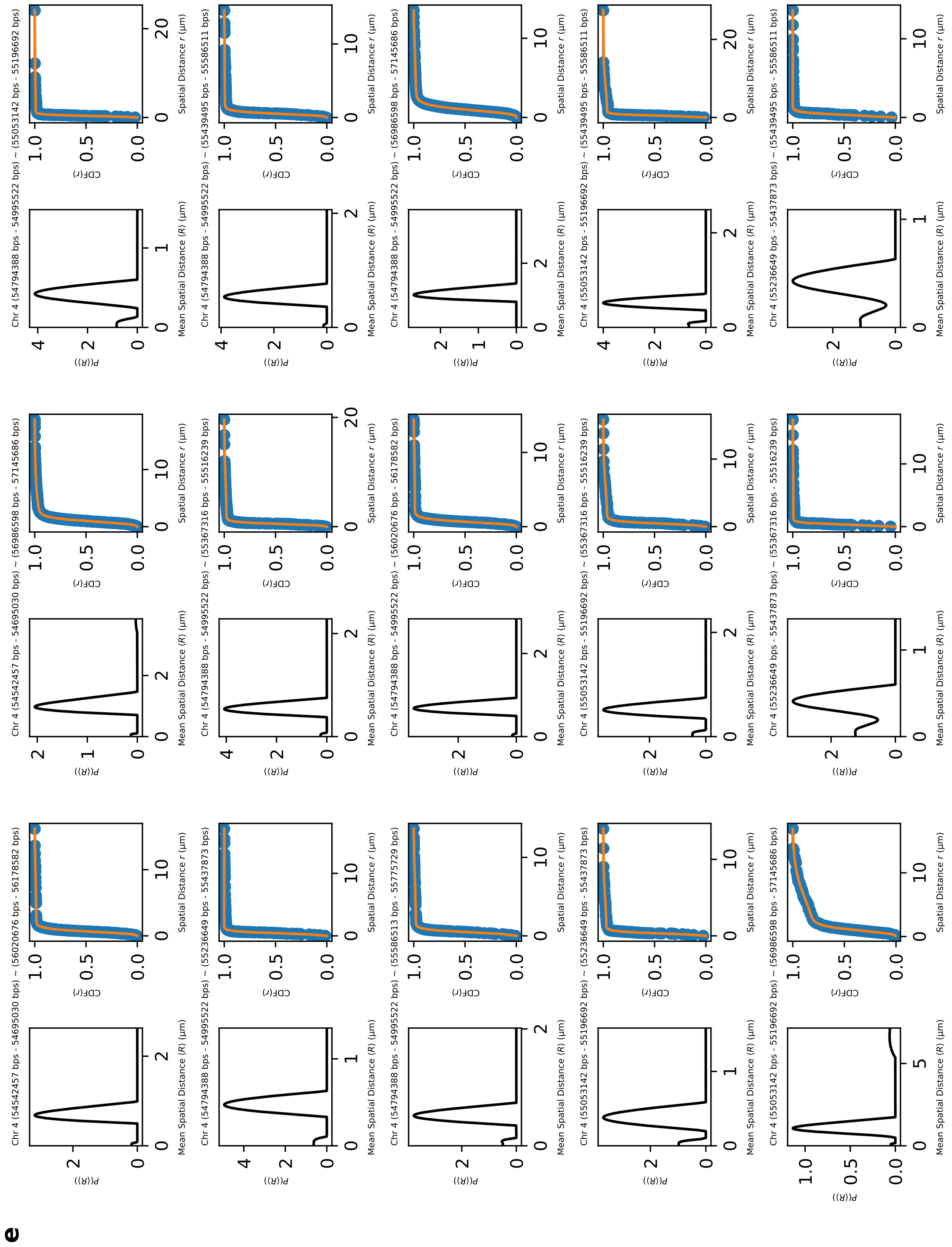
e
Chr 4 (54542457 bps - 54695030 bps) ~ (56020676 bps - 56178582 bps)
Chr 4 (54542457 bps - 54695030 bps) ~ (56986598 bps - 57145686 bps)
Chr 4 (54794388 bps - 54995522 bps) ~ (55053142 bps - 55196692 bps)
Chr 4 (54794388 bps - 54995522 bps) ~ (55236649 bps - 55437873 bps)
Chr 4 (54794388 bps - 54995522 bps) ~ (55367316 bps - 55516239 bps)
Chr 4 (54794388 bps - 54995522 bps) ~ (55439495 bps - 55586511 bps)
Chr 4 (54794388 bps - 54995522 bps) ~ (55586513 bps - 55775729 bps)
Chr 4 (54794388 bps - 54995522 bps) ~ (56020676 bps - 56178582 bps)
Chr 4 (54794388 bps - 54995522 bps) ~ (56986598 bps - 57145686 bps)
Chr 4 (55053142 bps - 55196692 bps) ~ (55236649 bps - 55437873 bps)
Chr 4 (55053142 bps - 55196692 bps) ~ (55367316 bps - 55516239 bps)
Chr 4 (55053142 bps - 55196692 bps) ~ (55439495 bps - 55586511 bps)
Chr 4 (55053142 bps - 55196692 bps) ~ (56986598 bps - 57145686 bps)
Chr 4 (55236649 bps - 55437873 bps) ~ (55367316 bps - 55516239 bps)
Chr 4 (55236649 bps - 55437873 bps) ~ (55439495 bps - 55586511 bps)
Mean Spatial Distance ⟨R⟩ (μm)
Spatial Distance r (μm)
P(⟨R⟩)
CDF(r)

**f**

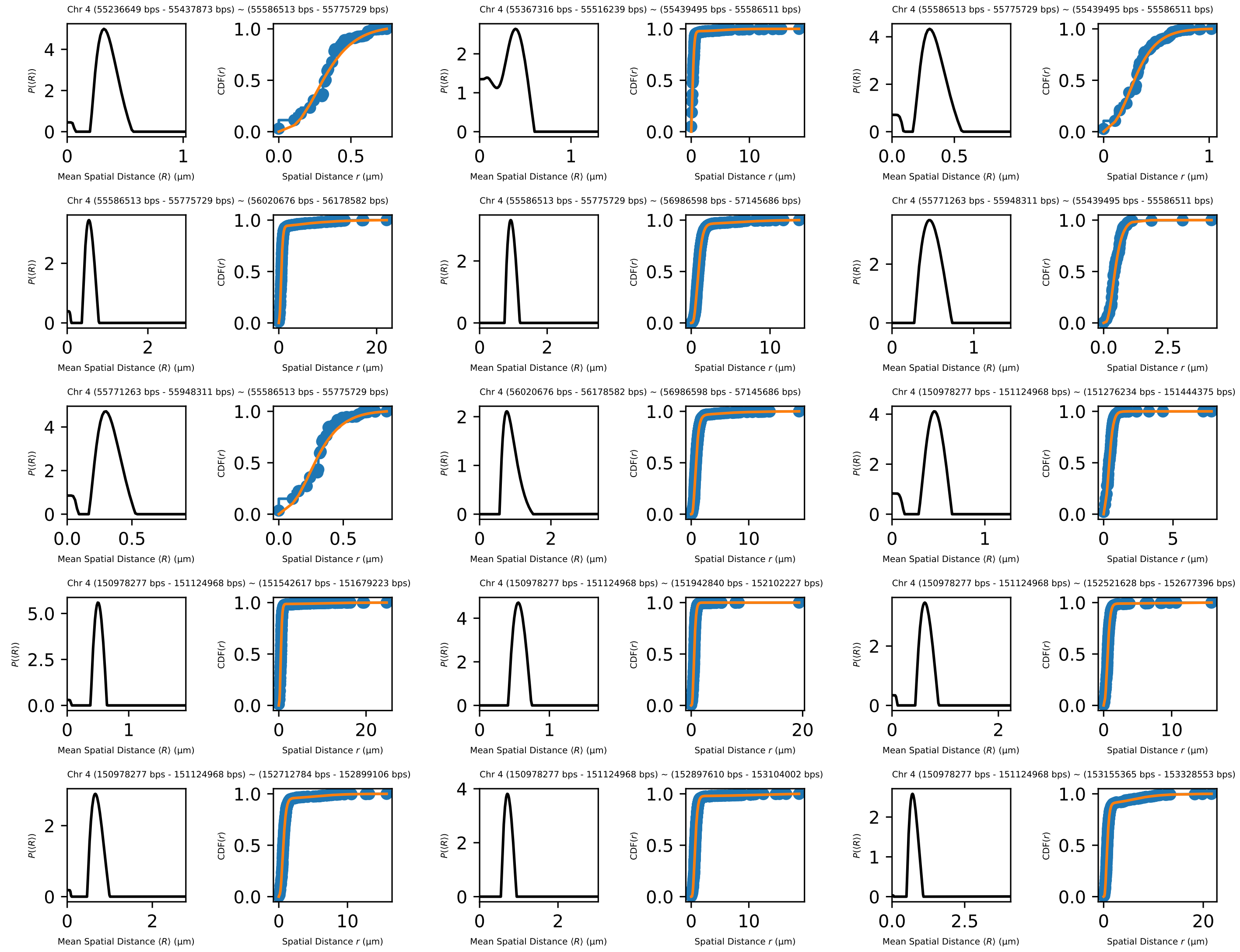
Chr 4 (55236649 bps - 55437873 bps) ~ (55586513 bps - 55775729 bps)
Chr 4 (55367316 bps - 55516239 bps) ~ (55439495 bps - 55586511 bps)
Chr 4 (55586513 bps - 55775729 bps) ~ (55439495 bps - 55586511 bps)
Chr 4 (55586513 bps - 55775729 bps) ~ (56020676 bps - 56178582 bps)
Chr 4 (55586513 bps - 55775729 bps) ~ (56986598 bps - 57145686 bps)
Chr 4 (55771263 bps - 55948311 bps) ~ (55439495 bps - 55586511 bps)
Chr 4 (55771263 bps - 55948311 bps) ~ (55586513 bps - 55775729 bps)
Chr 4 (56020676 bps - 56178582 bps) ~ (56986598 bps - 57145686 bps)
Chr 4 (150978277 bps - 151124968 bps) ~ (151276234 bps - 151444375 bps)
Chr 4 (150978277 bps - 151124968 bps) ~ (151542617 bps - 151679223 bps)
Chr 4 (150978277 bps - 151124968 bps) ~ (151942840 bps - 152102227 bps)
Chr 4 (150978277 bps - 151124968 bps) ~ (152521628 bps - 152677396 bps)
Chr 4 (150978277 bps - 151124968 bps) ~ (152712784 bps - 152899106 bps)
Chr 4 (150978277 bps - 151124968 bps) ~ (152897610 bps - 153104002 bps)
Chr 4 (150978277 bps - 151124968 bps) ~ (153155365 bps - 153328553 bps)
P(⟨R⟩)
Mean Spatial Distance ⟨R⟩ (μm)
CDF(r)
Spatial Distance r (μm)

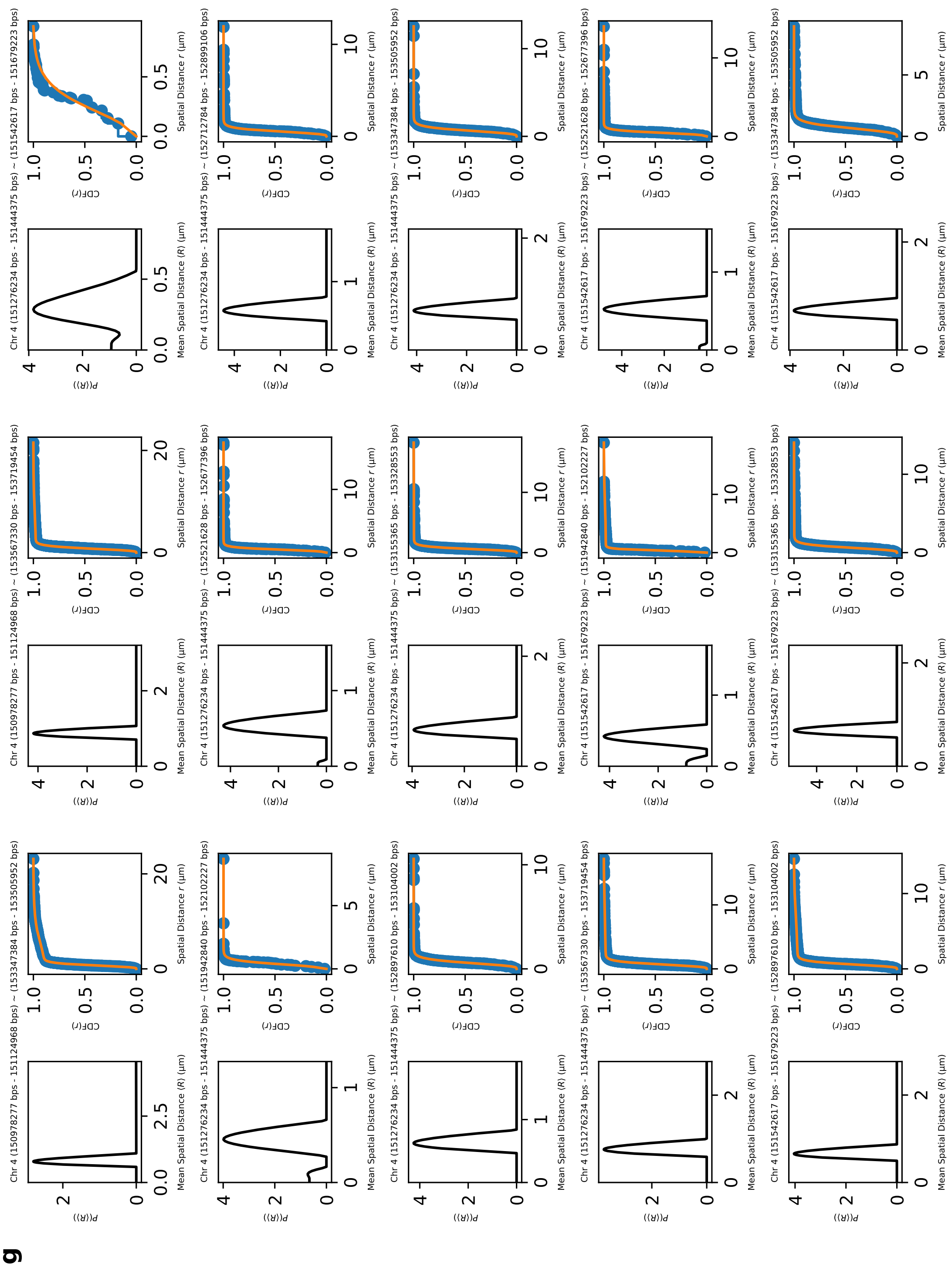
g
Chr 4 (150978277 bps - 151124968 bps) ~ (153347384 bps - 153505952 bps)
Chr 4 (150978277 bps - 151124968 bps) ~ (153567330 bps - 153719454 bps)
Chr 4 (151276234 bps - 151444375 bps) ~ (151542617 bps - 151679223 bps)
Chr 4 (151276234 bps - 151444375 bps) ~ (151942840 bps - 152102227 bps)
Chr 4 (151276234 bps - 151444375 bps) ~ (152521628 bps - 152677396 bps)
Chr 4 (151276234 bps - 151444375 bps) ~ (152712784 bps - 152899106 bps)
Chr 4 (151276234 bps - 151444375 bps) ~ (152897610 bps - 153104002 bps)
Chr 4 (151276234 bps - 151444375 bps) ~ (153155365 bps - 153328553 bps)
Chr 4 (151276234 bps - 151444375 bps) ~ (153347384 bps - 153505952 bps)
Chr 4 (151276234 bps - 151444375 bps) ~ (153567330 bps - 153719454 bps)
Chr 4 (151542617 bps - 151679223 bps) ~ (151942840 bps - 152102227 bps)
Chr 4 (151542617 bps - 151679223 bps) ~ (152521628 bps - 152677396 bps)
Chr 4 (151542617 bps - 151679223 bps) ~ (152897610 bps - 153104002 bps)
Chr 4 (151542617 bps - 151679223 bps) ~ (153155365 bps - 153328553 bps)
Chr 4 (151542617 bps - 151679223 bps) ~ (153347384 bps - 153505952 bps)
P(⟨R⟩)
Mean Spatial Distance ⟨R⟩ (μm)
CDF(r)
Spatial Distance r (μm)

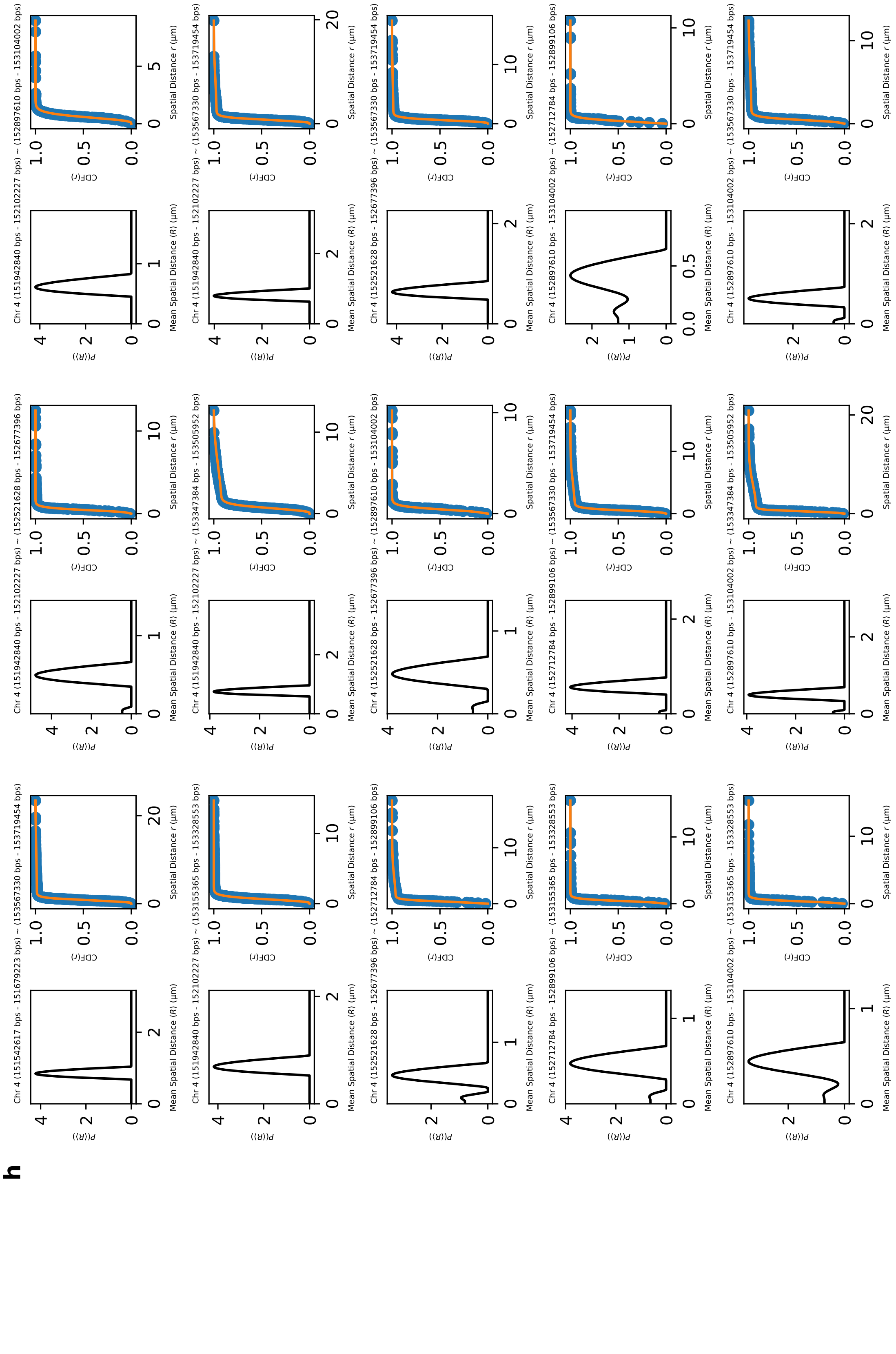
h

**i**

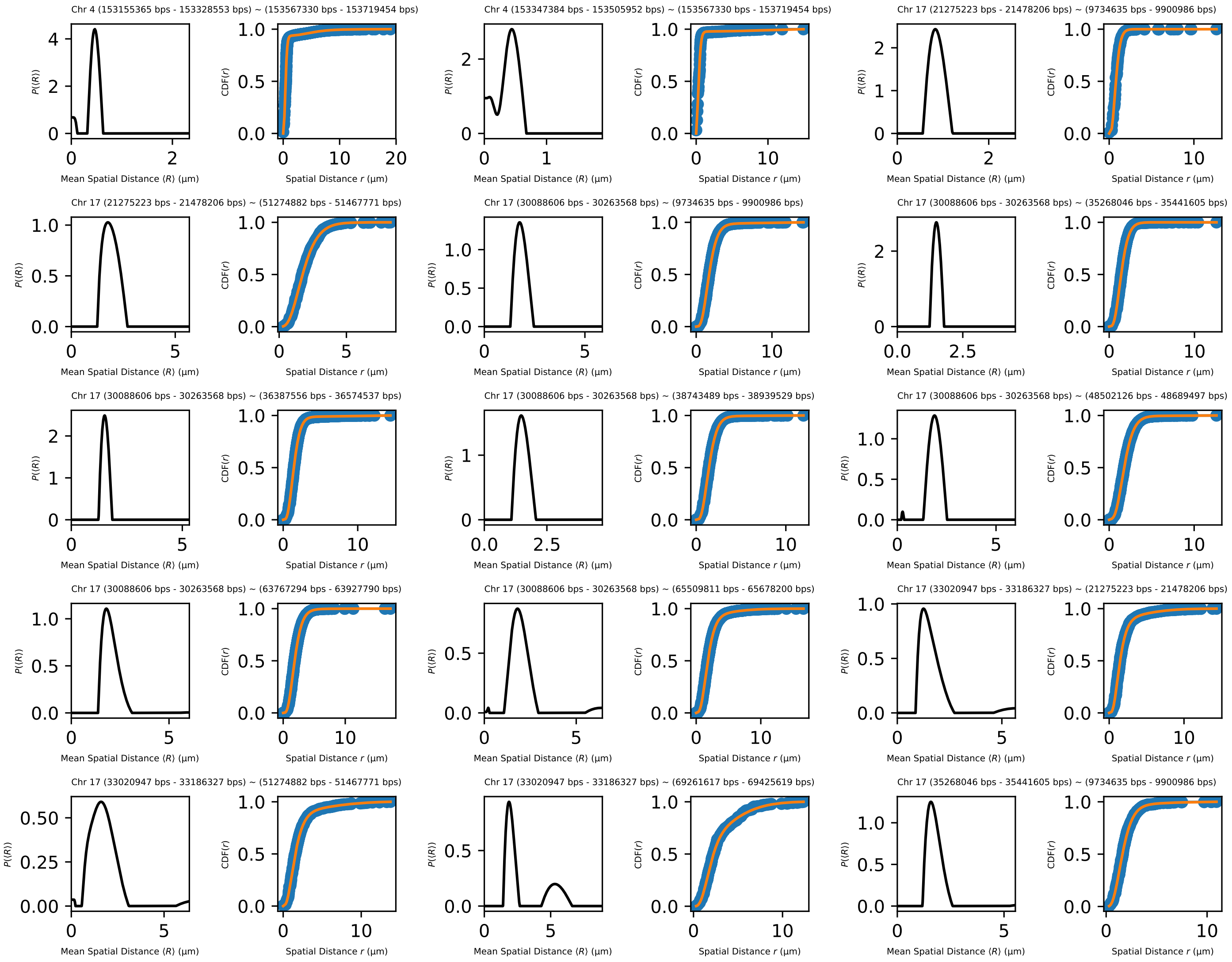
Chr 4 (153155365 bps - 153328553 bps) ~ (153567330 bps - 153719454 bps)
Chr 4 (153347384 bps - 153505952 bps) ~ (153567330 bps - 153719454 bps)
Chr 17 (21275223 bps - 21478206 bps) ~ (9734635 bps - 9900986 bps)
Chr 17 (21275223 bps - 21478206 bps) ~ (51274882 bps - 51467771 bps)
Chr 17 (30088606 bps - 30263568 bps) ~ (9734635 bps - 9900986 bps)
Chr 17 (30088606 bps - 30263568 bps) ~ (35268046 bps - 35441605 bps)
Chr 17 (30088606 bps - 30263568 bps) ~ (36387556 bps - 36574537 bps)
Chr 17 (30088606 bps - 30263568 bps) ~ (38743489 bps - 38939529 bps)
Chr 17 (30088606 bps - 30263568 bps) ~ (48502126 bps - 48689497 bps)
Chr 17 (30088606 bps - 30263568 bps) ~ (63767294 bps - 63927790 bps)
Chr 17 (30088606 bps - 30263568 bps) ~ (65509811 bps - 65678200 bps)
Chr 17 (33020947 bps - 33186327 bps) ~ (21275223 bps - 21478206 bps)
Chr 17 (33020947 bps - 33186327 bps) ~ (51274882 bps - 51467771 bps)
Chr 17 (33020947 bps - 33186327 bps) ~ (69261617 bps - 69425619 bps)
Chr 17 (35268046 bps - 35441605 bps) ~ (9734635 bps - 9900986 bps)
P(⟨R⟩)
CDF(r)
Mean Spatial Distance ⟨R⟩ (μm)
Spatial Distance r (μm)

**j**

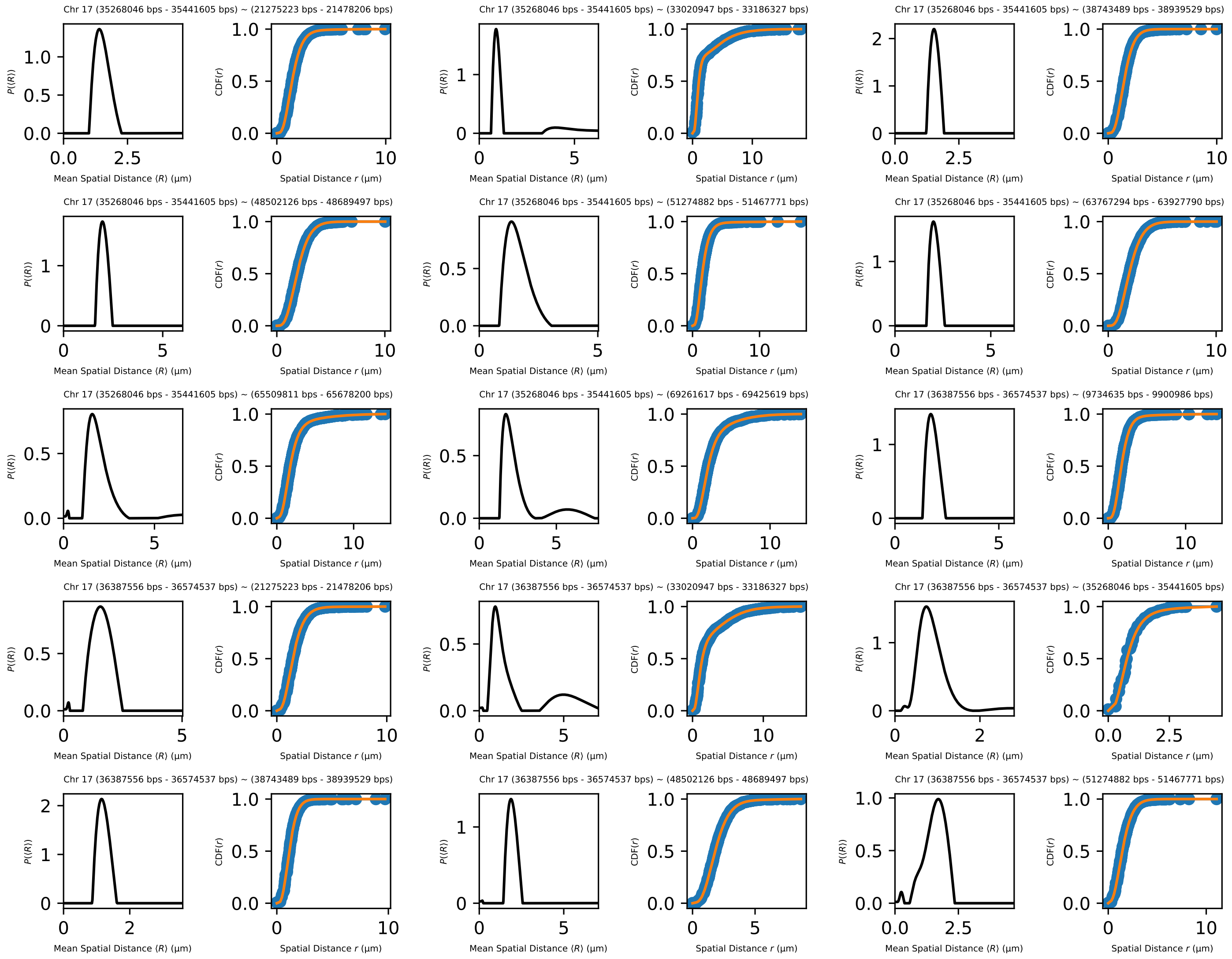
Chr 17 (35268046 bps - 35441605 bps) ~ (21275223 bps - 21478206 bps)
Chr 17 (35268046 bps - 35441605 bps) ~ (33020947 bps - 33186327 bps)
Chr 17 (35268046 bps - 35441605 bps) ~ (38743489 bps - 38939529 bps)
Chr 17 (35268046 bps - 35441605 bps) ~ (48502126 bps - 48689497 bps)
Chr 17 (35268046 bps - 35441605 bps) ~ (51274882 bps - 51467771 bps)
Chr 17 (35268046 bps - 35441605 bps) ~ (63767294 bps - 63927790 bps)
Chr 17 (35268046 bps - 35441605 bps) ~ (65509811 bps - 65678200 bps)
Chr 17 (35268046 bps - 35441605 bps) ~ (69261617 bps - 69425619 bps)
Chr 17 (36387556 bps - 36574537 bps) ~ (9734635 bps - 9900986 bps)
Chr 17 (36387556 bps - 36574537 bps) ~ (21275223 bps - 21478206 bps)
Chr 17 (36387556 bps - 36574537 bps) ~ (33020947 bps - 33186327 bps)
Chr 17 (36387556 bps - 36574537 bps) ~ (35268046 bps - 35441605 bps)
Chr 17 (36387556 bps - 36574537 bps) ~ (38743489 bps - 38939529 bps)
Chr 17 (36387556 bps - 36574537 bps) ~ (48502126 bps - 48689497 bps)
Chr 17 (36387556 bps - 36574537 bps) ~ (51274882 bps - 51467771 bps)
P(⟨R⟩)
CDF(r)
Mean Spatial Distance ⟨R⟩ (μm)
Spatial Distance r (μm)

**k**

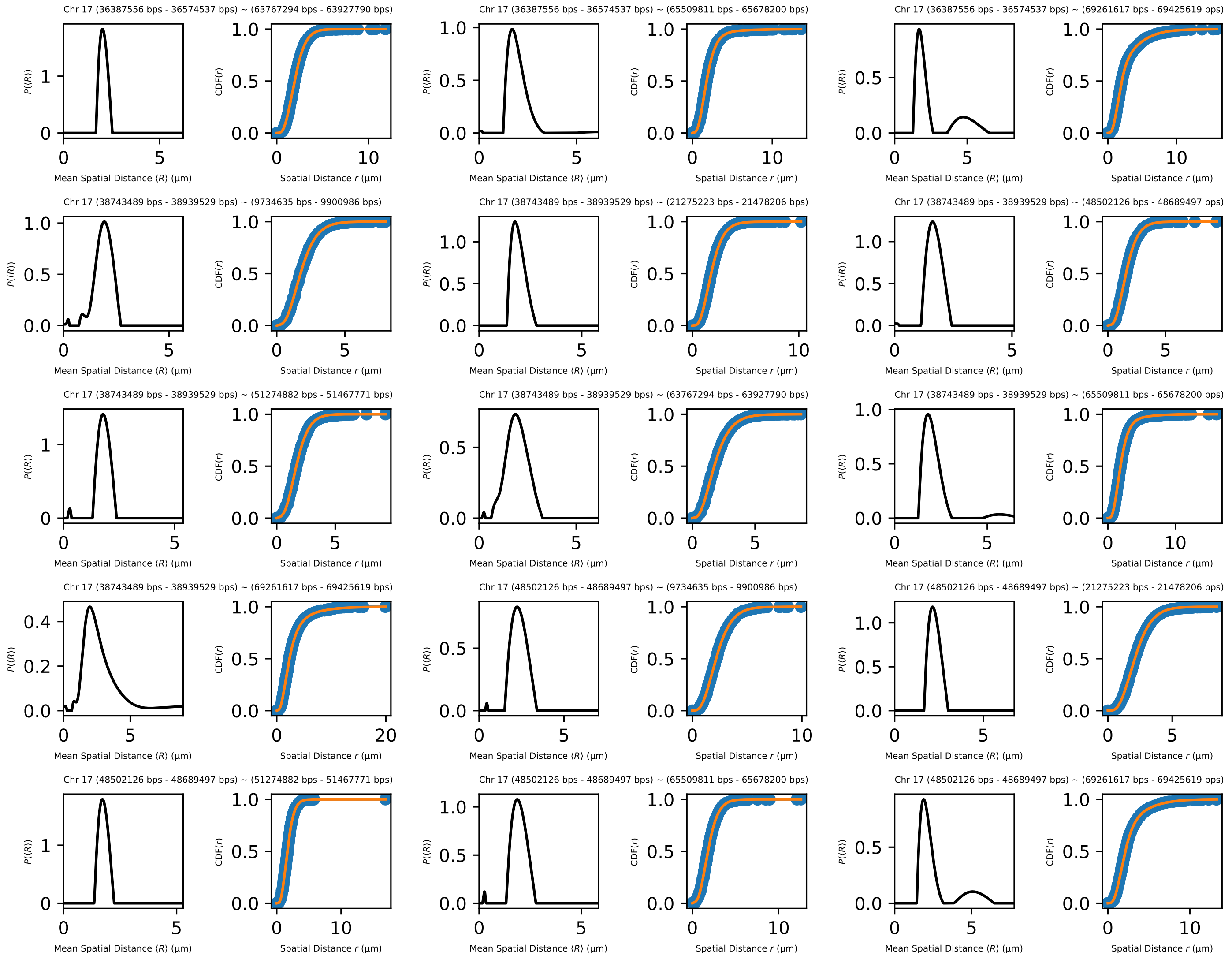

Chr 17 (36387556 bps - 36574537 bps) ~ (63767294 bps - 63927790 bps)
Chr 17 (36387556 bps - 36574537 bps) ~ (65509811 bps - 65678200 bps)
Chr 17 (36387556 bps - 36574537 bps) ~ (69261617 bps - 69425619 bps)
Chr 17 (38743489 bps - 38939529 bps) ~ (9734635 bps - 9900986 bps)
Chr 17 (38743489 bps - 38939529 bps) ~ (21275223 bps - 21478206 bps)
Chr 17 (38743489 bps - 38939529 bps) ~ (48502126 bps - 48689497 bps)
Chr 17 (38743489 bps - 38939529 bps) ~ (51274882 bps - 51467771 bps)
Chr 17 (38743489 bps - 38939529 bps) ~ (63767294 bps - 63927790 bps)
Chr 17 (38743489 bps - 38939529 bps) ~ (65509811 bps - 65678200 bps)
Chr 17 (38743489 bps - 38939529 bps) ~ (69261617 bps - 69425619 bps)
Chr 17 (48502126 bps - 48689497 bps) ~ (9734635 bps - 9900986 bps)
Chr 17 (48502126 bps - 48689497 bps) ~ (21275223 bps - 21478206 bps)
Chr 17 (48502126 bps - 48689497 bps) ~ (51274882 bps - 51467771 bps)
Chr 17 (48502126 bps - 48689497 bps) ~ (65509811 bps - 65678200 bps)
Chr 17 (48502126 bps - 48689497 bps) ~ (69261617 bps - 69425619 bps)
Mean Spatial Distance ⟨R⟩ (µm)
Spatial Distance r (µm)
P(⟨R⟩)
CDF(r)

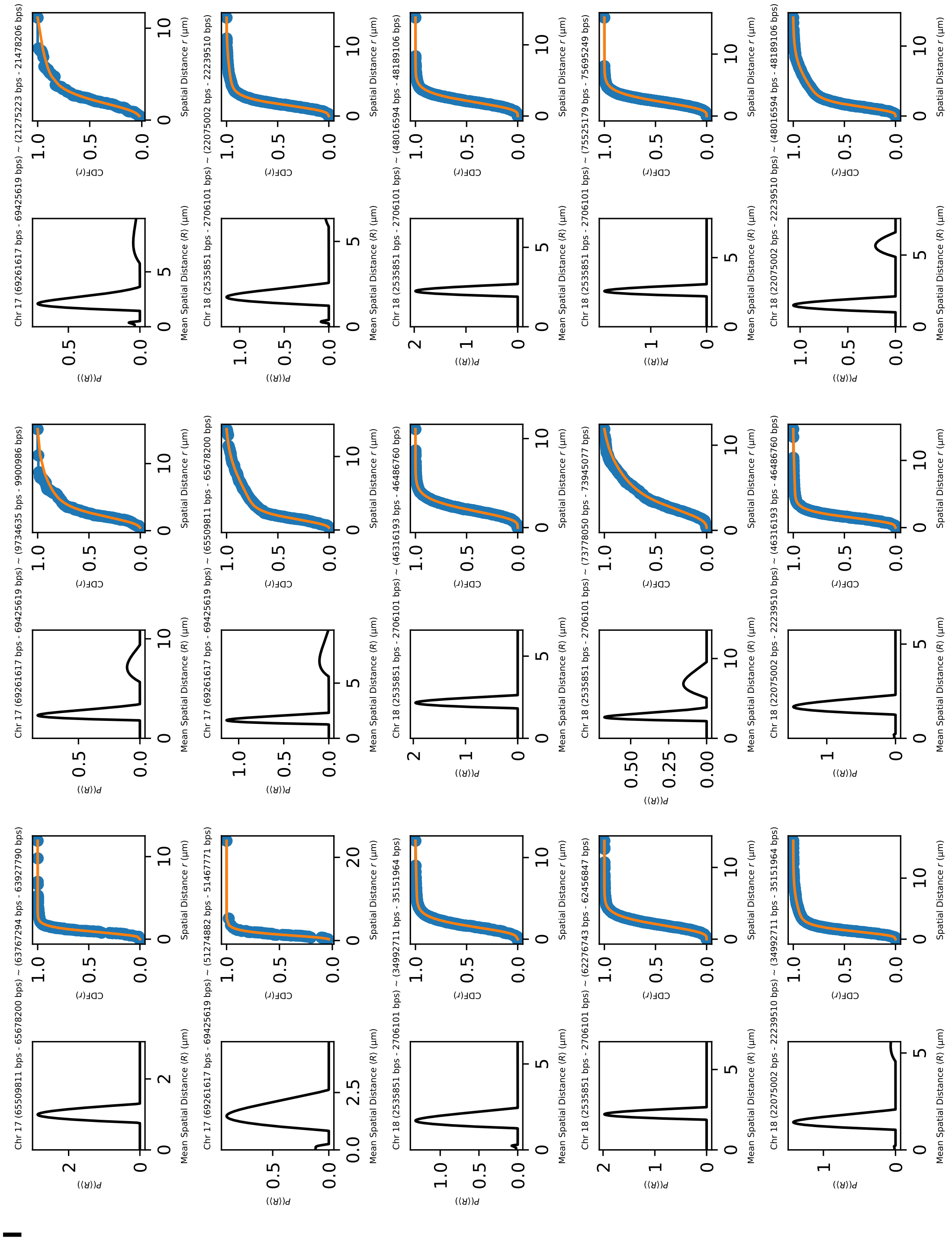

Chr 17 (65509811 bps - 65678200 bps) ~ (63767294 bps - 63927790 bps)
Chr 17 (69261617 bps - 69425619 bps) ~ (9734635 bps - 9900986 bps)
Chr 17 (69261617 bps - 69425619 bps) ~ (21275223 bps - 21478206 bps)
Chr 17 (69261617 bps - 69425619 bps) ~ (51274882 bps - 51467771 bps)
Chr 17 (69261617 bps - 69425619 bps) ~ (65509811 bps - 65678200 bps)
Chr 18 (2535851 bps - 2706101 bps) ~ (22075002 bps - 22239510 bps)
Chr 18 (2535851 bps - 2706101 bps) ~ (34992711 bps - 35151964 bps)
Chr 18 (2535851 bps - 2706101 bps) ~ (46316193 bps - 46486760 bps)
Chr 18 (2535851 bps - 2706101 bps) ~ (48016594 bps - 48189106 bps)
Chr 18 (2535851 bps - 2706101 bps) ~ (62276743 bps - 62456847 bps)
Chr 18 (2535851 bps - 2706101 bps) ~ (73778050 bps - 73945077 bps)
Chr 18 (2535851 bps - 2706101 bps) ~ (75525179 bps - 75695249 bps)
Chr 18 (22075002 bps - 22239510 bps) ~ (34992711 bps - 35151964 bps)
Chr 18 (22075002 bps - 22239510 bps) ~ (46316193 bps - 46486760 bps)
Chr 18 (22075002 bps - 22239510 bps) ~ (48016594 bps - 48189106 bps)
Mean Spatial Distance ⟨R⟩ (μm)
Spatial Distance r (μm)
P(⟨R⟩)
CDF(r)

**m**

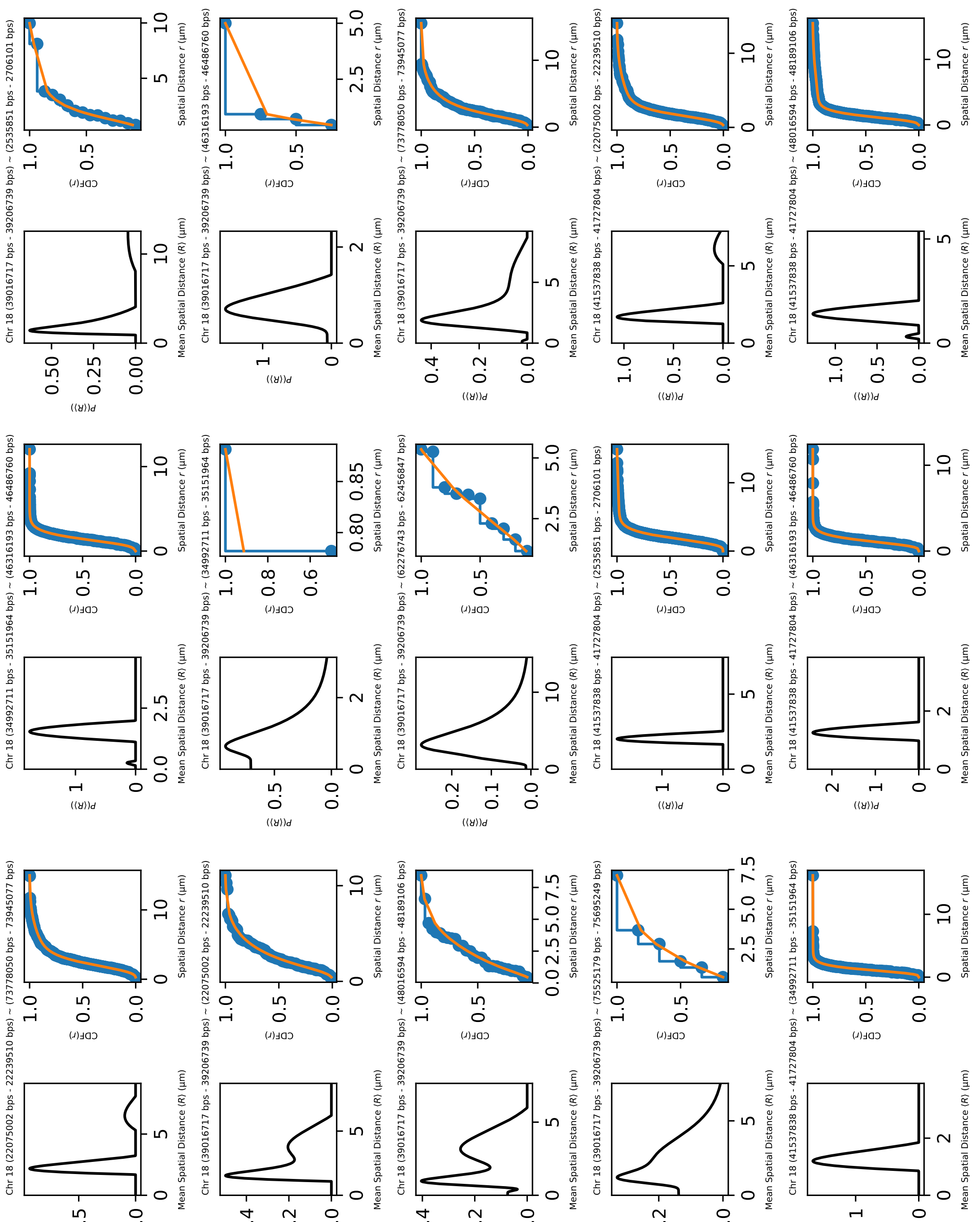

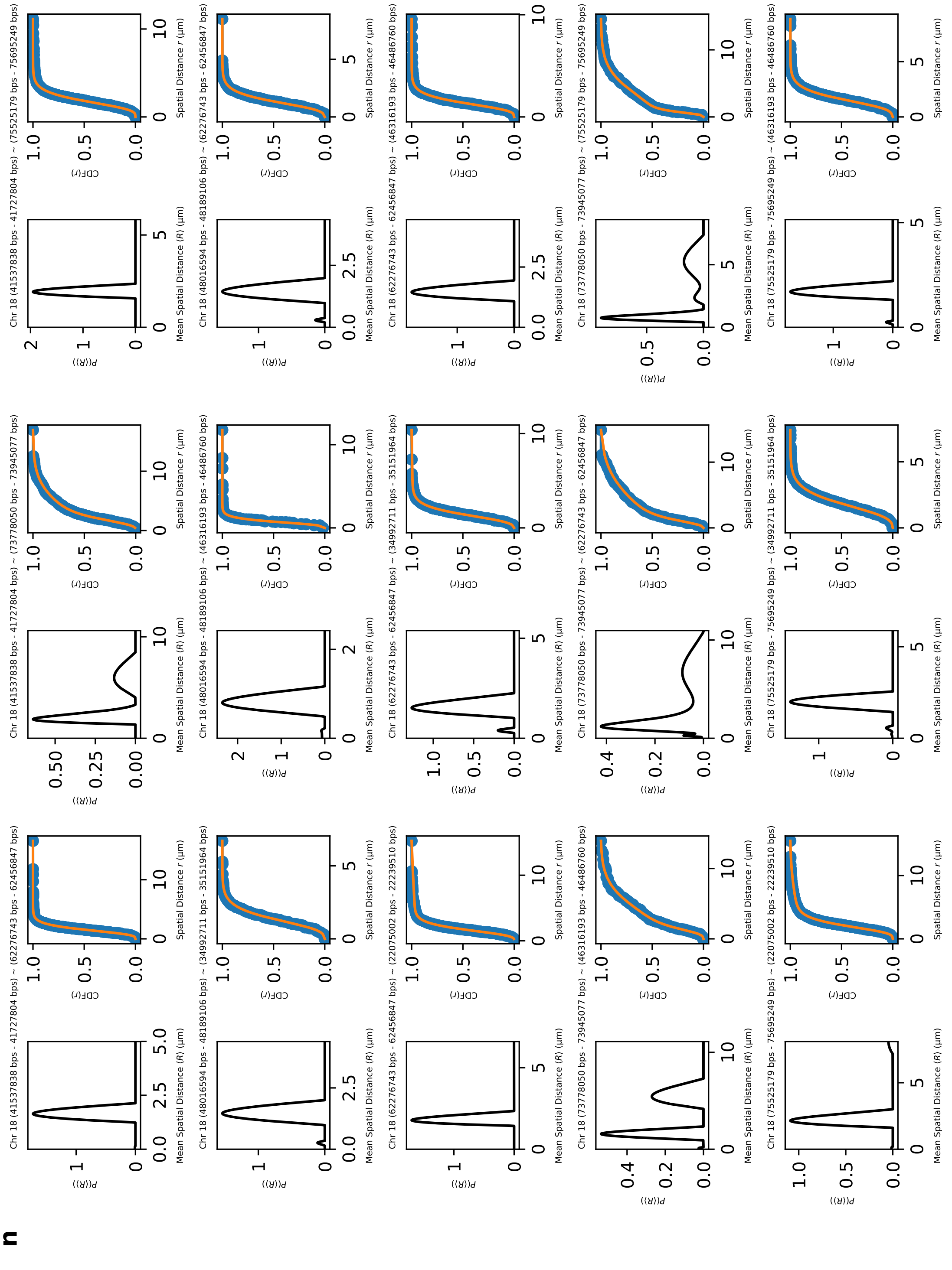
n
Chr 18 (41537838 bps - 41727804 bps) ~ (62276743 bps - 62456847 bps)
Chr 18 (41537838 bps - 41727804 bps) ~ (73778050 bps - 73945077 bps)
Chr 18 (41537838 bps - 41727804 bps) ~ (75525179 bps - 75695249 bps)
Chr 18 (48016594 bps - 48189106 bps) ~ (34992711 bps - 35151964 bps)
Chr 18 (48016594 bps - 48189106 bps) ~ (46316193 bps - 46486760 bps)
Chr 18 (48016594 bps - 48189106 bps) ~ (62276743 bps - 62456847 bps)
Chr 18 (62276743 bps - 62456847 bps) ~ (22075002 bps - 22239510 bps)
Chr 18 (62276743 bps - 62456847 bps) ~ (34992711 bps - 35151964 bps)
Chr 18 (62276743 bps - 62456847 bps) ~ (46316193 bps - 46486760 bps)
Chr 18 (73778050 bps - 73945077 bps) ~ (46316193 bps - 46486760 bps)
Chr 18 (73778050 bps - 73945077 bps) ~ (62276743 bps - 62456847 bps)
Chr 18 (73778050 bps - 73945077 bps) ~ (75525179 bps - 75695249 bps)
Chr 18 (75525179 bps - 75695249 bps) ~ (22075002 bps - 22239510 bps)
Chr 18 (75525179 bps - 75695249 bps) ~ (34992711 bps - 35151964 bps)
Chr 18 (75525179 bps - 75695249 bps) ~ (46316193 bps - 46486760 bps)
Mean Spatial Distance ⟨R⟩ (μm)
Spatial Distance r (μm)

**o**

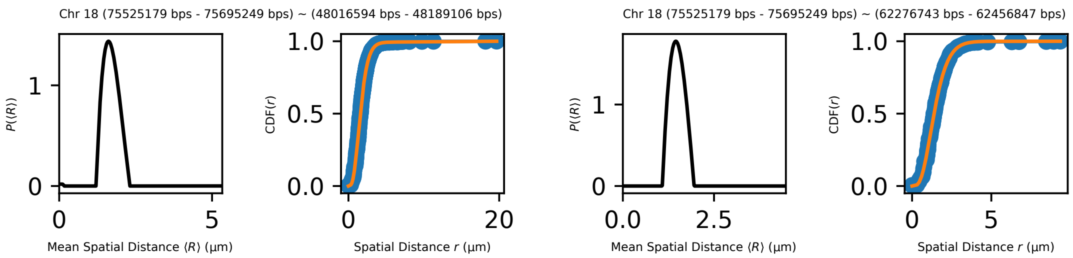


Supplementary Figure 8: (**a-o**) The fits of CDF($r$) using Supplementary Equation 9 to a total number of 212 pairs [5]. First 14 pages each show 15 pairs of loci and the last page shows 2 pairs of loci. For almost all the 212 pairs, orange lines, showing the fits using our theory, are indistinguishable from the experiment (blue markers). The distribution $P(\langle R\rangle)$ is solved using non-negative Tikhonov Regularization (Supplementary Note 7).
